# Biotic and abiotic factors shaping the genome of cockle (*Cerastoderma edule*) in the Northeast Atlantic: a baseline for sustainable management of its wild resources

**DOI:** 10.1101/2020.12.17.423063

**Authors:** Manuel Vera, Francesco Maroso, Sophie-B. Wilmes, Miguel Hermida, Andrés Blanco, Carlos Fernández, Emily Groves, Shelagh K Malham, Carmen Bouza, The Cockle’s Consortium, Peter E. Robins, Paulino Martínez

**Affiliations:** Department of Zoology, Genetics and Physical Anthropology. ACUIGEN group. Faculty of Veterinary. Universidade de Santiago de Compostela. Campus of Lugo. 27002 Lugo, Spain; Institute of Aquaculture, Universidade de Santiago de Compostela, 15705 Santiago de Compostela, Spain; Department of Life Sciences and Biotechnologies, University of Ferrara, via L. Borsari 46, 44124 Ferrara, Italy; School of Ocean Sciences, Marine Centre Wales, Bangor University, Menai Bridge, UK

**Author notes:** M. Vera, F. Maroso and S.-B. Wilmes have contributed equally to this work and are first co-authors.

**Keywords:** 2b-RAD, genetic structure, adaptive variation, larval dispersion modelling, fisheries management

## Abstract

Knowledge on how environmental factors shape the genome of marine species is crucial for sustainable management of fisheries and wild populations. The edible cockle (*Cerastoderma edule*) is a marine bivalve distributed along the Northeast Atlantic coast of Europe and is an important resource from both commercial and ecological perspectives. We performed a population genomics screening using 2b-RAD genotyping on 9,309 SNPs localised in the cockle’s genome on a sample of 536 specimens pertaining to 14 beds in the Northeast Atlantic to ascertain its genetic structure regarding environmental variation. Larval dispersal modelling considering species behaviour and interannual variability in ocean conditions was carried out, as an essential background to compare genetic information with. Cockle populations in the Northeast Atlantic were shown to be panmictic and displayed low but significant geographical differentiation across populations (F_ST_ = 0.0240; P < 0.001), albeit not across generations. We identified 441 outlier SNPs related to divergent selection, sea surface temperature being the main environmental driver following a latitudinal axis. Two main genetic groups were identified, northwards and southwards of French Brittany, in accordance with our modelling, which demonstrated a barrier for larval dispersal linked to the Ushant front. Further genetic subdivision was observed using outlier loci and considering larval behaviour. The northern group was divided into the Irish/Celtic Seas and the English Channel/North Sea, while the southern group was divided into three subgroups. This information represents the baseline for management of cockles, designing conservation strategies, founding broodstock for depleted beds, and producing suitable seed for aquaculture production.

## Introduction

The genetic structure of marine species and patterns of connectivity between discrete populations are central to their health and resilience to external pressures such as parasites and pathogens (Rowley et al., 2014), pollution, sustainable management and exploitation, and climate change over ecological and evolutionary timescales (Cowen & Sponaugle, 2009; Burgess et al., 2014). Detection of genetic structure in marine species remains challenging due to the often large effective population sizes and high levels of gene flow facilitated by the scarcity of physical barriers, which lead to genomic homogenization (Danancher & Garcia-Vazquez, 2011; do Prado et al., 2018; Zhao et al., 2018). However, genetic differentiation can be enhanced by oceanic features such as current systems, fronts, gyres and eddies (Nielsen et al., 2004; Blanco-González et al., 2016; Vera et al., 2016; Xuereb et al., 2018), and natural selection in response to environmental variation (Vilas et al., 2015; Clucas et al., 2019; Jiménez-Mena et al., 2020). Distinguishing between neutral and adaptive genetic variation in the marine landscape has become a central issue in conservation biology, allowing for interpreting genetic variation in both historical/demographic and adaptive terms (Nielsen et al., 2009; Bernatchez, 2016). This information is essential for identifying broodstock with the necessary genetic diversity for conservation and breeding programs in marine aquaculture (do Prado et al., 2018; Hughes et al., 2019).

The edible cockle, *Cerastoderma edule*, is a bivalve mollusc naturally distributed along the Northeast Atlantic coast, from Senegal to Norway, inhabiting intertidal soft sediments (Hayward & Ryland, 1995). The species plays a crucial role as a food source for birds, crustaceans and fish (Norris et al., 1998). Moreover, the species is highly appreciated for cuisine, and its main fisheries are located in Ireland, the United Kingdom, France, Spain and Portugal, where its commercialisation represents employment of thousands of collectors, processors and sellers (http://www.cockles-project.eu/). *C*. *edule* is dioecious and can live up to ten years with a fast sexual maturation (reached in its first year) and high fecundity (Honkoop & van der Meer, 1998), with reproductive periods spanning from late spring to mid-autumn (Malham et al., 2012; Mahony et al., 2020). Larvae are planktonic and remain in the water column for around 30 days, which allows for passive larval dispersal by ocean currents that drive connectivity and gene flow between populations spread along the Northeast Atlantic coast (de Montaudouin, et al., 2003; Dare et al., 2004).

The genetic structure of *C*. *edule* across the Northeast Atlantic has been studied over the last forty years. Pioneering studies using allozymes detected genetic differences between populations located on either side of the English Channel (collected in Wales, France and The Netherlands; Beaumont et al., 1980), but also a high connectivity and gene flow from France to Denmark (Hummel et al., 1994). Mitochondrial DNA (mtDNA) sequencing carried out on wider sampling (from Morocco to Russia) revealed the presence of two major mtDNA groups in northern and southern areas, suggesting the presence of a northern cryptic refugia for *C. edule* (Krakau et al., 2012; Martínez et al., 2015). Studies using microsatellites showed high homogeneity in the southern beds from Portugal and Spain (Martínez et al., 2013; 2015), while two main clusters were identified in the northern area in the British Isles and the North Sea. In summary, three major areas were defined from microsatellite data: (*i*) a southern region (Morocco, Portugal, Spain and French beds to the English Channel); (*ii*) Ireland, Great Britain and southern North Sea (The Netherlands and Germany); (*iii*) a northern group (Scotland, Denmark, Norway and Russia) (Martínez et al., 2015). However, the low amount of microsatellite markers used have greatly limited the investigation on local adaptation and population connectivity at the fine scale necessary for the appropriate management of the exploited species (Bernatchez et al., 2017).

Recently, Coscia et al. (2020) analysed the genetic structure and connectivity among cockle populations within the Celtic/Irish seas using data from Restriction-site Associated DNA sequencing (RADseq) and a population genomics approach in combination with information on ocean conditions and larval dispersal modelling. They identified a significant genetic differentiation in a small geographical area (three groups; F_ST_ = 0.021) mainly driven by salinity, larval dispersal by oceanographic currents and geographical distance. These results show that a finer structure underlies cockle distribution in the Northeast Atlantic and that a genomic scan covering southern and northern beds taking as background the dispersal of larvae in the area is necessary to understand how the species is structured for its appropriate management.

In this study, we applied a 2b-RADseq genotyping by sequencing approach to assess the genetic structure of *C. edule* along the Northeaster Atlantic coast considering environmental drivers and larval dispersion models data with the aim of providing essential information for the sustainable management of its resources. Major regions previously identified with microsatellites were confirmed, but refined information was obtained, mostly in agreement with the ocean dynamics and resulting larval dispersal patterns across the Northeast Atlantic.

## Material and methods

### Oceanography of the study area

The study area covers the Northeast Atlantic from southeast Portugal to northeast Ireland and the southern North Sea (Fig. 1). This area is divided into several oceanographic regions: Iberian coastal waters, the Bay of Biscay, the English Channel, the Celtic Sea, the Irish Sea and the North Sea. These regions are to some extent discrete units with limited oceanographic connectivity between them resulting from either divergent coastal currents or frontal currents generated during summer heating (Galparoso et al., 2014). During summer months, when upwelling is a prominent feature along the Galician coast (NW Spain), the Portuguese coastal currents transports surface waters southward along the Galician and Portuguese west coast, whereas during the winter months the Iberian Poleward Current shoals and moves surface waters northwards (Teles-Machado et al., 2016). Along the northern coast of Spain, a strong westward transport develops during the summer months which changes direction during the winter months and links into the slope current along the Amorican and Aquitaine Shelf (W France). The coastal circulation along the French coast of the Bay of Biscay is characterised by northward transport by the Iberian Poleward Current during the winter which reverses in direction and reduces in strength during the summer months (Charria et al., 2013). Tidal mixing fronts separating mixed from seasonally stratified waters form in early summer and extend into the autumn at the entrance to the English Channel (Ushant Front) (Sournia et al., 1988) and between southwest Wales and southeast Ireland (Celtic Sea Front). From late spring to early autumn a current system develops in the Celtic Sea that transports waters from southwestern Britain via the frontal jet associated with the Celtic Sea Front to the south and west coasts of Ireland as the Irish coastal current (Horsburgh et al., 1998; Brown et al., 2003; Fernand et al., 2006).

**Figure 1.**
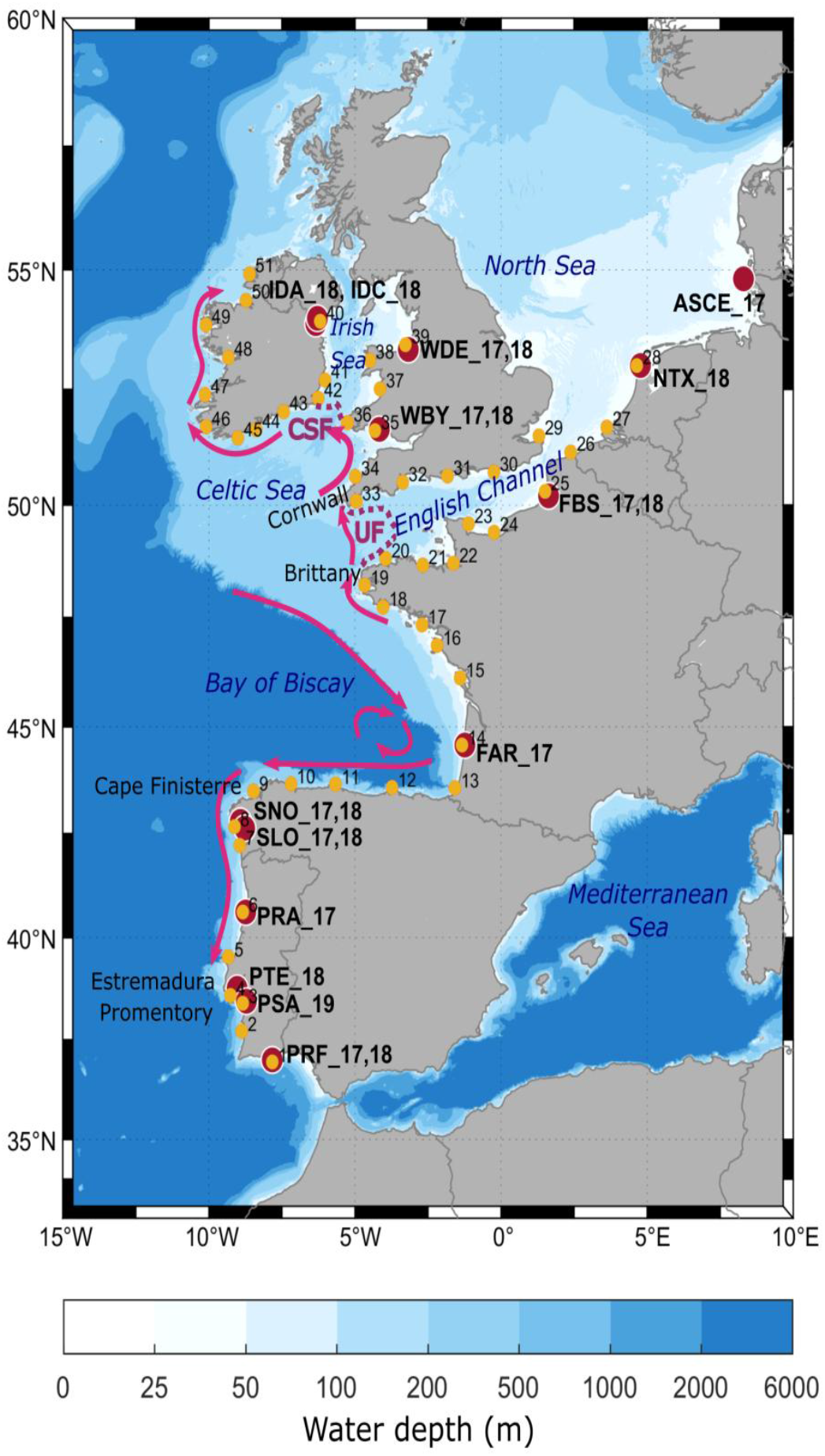
Study area for *C*. *edule* genetic analysis and larval dispersal modelling. Ocean bathymetry is shaded in blue. Summer surface currents are schematically represented by magenta-coloured arrows (see section Study Area for a detailed description and references). Locations of fronts are depicted by purple dotted lines (CSF = Celtic Sea Front; UF = Ushant Front). Location of the *C*. *edule* beds for the genetic analysis are shown in dark red (see Table 1 for location codes) and particle release locations for larval dispersal modeling are shown in yellow and numbered from 1 to 51. Location codes in panel A are detailed in Table 1.

### Sampling

A total of 545 cockles from 14 natural beds distributed along the Northeast Atlantic Coast (Fig. 1) were collected during the period 2017-2019 and stored in 100% ethanol (Table 1). Temporal replicates, to analyse genetic stability across generations, were obtained for six beds in 2017 and 2018. Specimens collected belonged to the 0+ year age class to avoid generation overlapping between consecutive year cohorts.

**Table 1.** Sampling and genetic diversity features of the edible cockle (*Cerastoderma edule*) beds studied in the Atlantic Area

| Bed | Sampling year | Latitude (deg N) | Longitude (deg E) | Code | Country | N | Complete dataset |  |  | Polymorphic loci (MAF > 0.010) |  |  |
| --- | --- | --- | --- | --- | --- | --- | --- | --- | --- | --- | --- | --- |
|  |  |  |  |  |  |  | Ho | He | F<sub>IS</sub> | Ho | He | F<sub>IS</sub> |
| Sylt | 2017 | 54.814 | 8.298 | ASCE_17 | Germany | 22 (22) | 0.070 | 0.077 | 0.089 | 0.172 | 0.191 | 0.099 |
| Texel | 2018 | 53.004 | 4.771 | NTX_18 | Netherlands | 21 (21) | 0.075 | 0.078 | 0.034 | 0.183 | 0.193 | 0.052 |
| Dee Estuary | 2017 | 53.343 | -3.174 | WDE_17 | Wales | 30 (28) | 0.071 | 0.078 | 0.079 | 0.165 | 0.182 | 0.093 |
| Dee Estuary | 2018 | 53.343 | -3.174 | WDE_18 | Wales | 30 (30) | 0.075 | 0.083 | 0.099 | 0.153 | 0.172 | 0.110 |
| Burry | 2017 | 51.643 | -4.166 | WBY_17 | Wales | 30 (30) | 0.073 | 0.081 | 0.092 | 0.148 | 0.165 | 0.103 |
| Burry | 2018 | 51.643 | -4.166 | WBY_18 | Wales | 30 (30) | 0.070 | 0.077 | 0.098 | 0.157 | 0.176 | 0.108 |
| Dundalk Bay-Annagassan | 2018 | 53.884 | -6.341 | IDA_18 | Ireland | 29 (29) | 0.074 | 0.081 | 0.083 | 0.159 | 0.176 | 0.097 |
| Dundalk Bay-Cooley | 2018 | 53.996 | -6.287 | IDC_18 | Ireland | 22 (22) | 0.077 | 0.084 | 0.084 | 0.173 | 0.190 | 0.089 |
| Somme Bay | 2017 | 50.201 | 1.627 | FBS_17 | France | 30 (30) | 0.072 | 0.080 | 0.110 | 0.146 | 0.166 | 0.120 |
| Somme Bay | 2018 | 50.201 | 1.627 | FBS_18 | France | 31 (31) | 0.072 | 0.079 | 0.084 | 0.149 | 0.165 | 0.097 |
| Archachon Bay | 2017 | 44.580 | -1.238 | FAR_17 | France | 30 (30) | 0.074 | 0.083 | 0.111 | 0.138 | 0.157 | 0.121 |
| Ría de Noia | 2017 | 42.790 | -8.923 | SNO_17 | Spain | 30 (30) | 0.079 | 0.088 | 0.098 | 0.139 | 0.155 | 0.103 |
| Ría de Noia | 2018 | 42.790 | -8.923 | SNO_18 | Spain | 30(30) | 0.072 | 0.081 | 0.118 | 0.137 | 0.158 | 0.133 |
| Lombos do Ulla | 2017 | 42.629 | -8.775 | SLO_17 | Spain | 30 (30) | 0.076 | 0.086 | 0.114 | 0.136 | 0.155 | 0.123 |
| Lombos do Ulla | 2018 | 42.629 | -8.775 | SLO_18 | Spain | 32 (32) | 0.073 | 0.083 | 0.119 | 0.134 | 0.153 | 0.124 |
| Ria de Aveiro | 2017 | 40.623 | -8.739 | PRA_17 | Portugal | 30 (30) | 0.072 | 0.081 | 0.113 | 0.141 | 0.161 | 0.124 |
| Tejo Estuary | 2018 | 38.767 | -9.033 | PTE_18 | Portugal | 8 (8) | 0.073 | 0.078 | 0.063 | 0.271 | 0.290 | 0.066 |
| Sado Estuary | 2019 | 38.450 | -8.717 | PSA_19 | Portugal | 20 (20) | 0.071 | 0.080 | 0.108 | 0.177 | 0.202 | 0.124 |
| Ria Formosa | 2017 | 36.998 | -7.830 | PRF_17 | Portugal | 30 (30) | 0.079 | 0.084 | 0.066 | 0.146 | 0.158 | 0.076 |
| Ria Formosa | 2018 | 36.998 | -7.830 | PRF_18 | Portugal | 30 (23) | 0.079 | 0.086 | 0.079 | 0.159 | 0.174 | 0.086 |

N: number of individuals analysed (in parentheses N after quality filtering); Ho: observed heterozytosity; He: expected heterozygosity; Fis: intrapopulation fixation index.

### Single Nucleotide Polymorphism (SNP) genotyping

Total DNA was extracted from gill tissue samples using the e.Z.N.A. E-96 mollusc DNA kit (OMEGA Bio-tech), following manufacturer recommendations. SNP identification and selection, as well as genotyping and validation protocols followed those described by Maroso et al. (2019). Briefly, AlfI IIb restriction enzyme (RE) was used to construct the 2b-RAD libraries, which were evenly pooled for sequencing in Illumina Next-seq including 90 individuals per run. The recently assembled cockle’s genome (Tubío et al., unpublished data) was used to align reads from each individual using Bowtie 1.1.2 (Langmead et al., 2009), allowing a maximum of three mismatches and a unique valid alignment (-v 3 -m 1). Stacks 2.0 (Catchen et al., 2013) was then used to call SNPs and genotype a common set of markers in the sample set, applying the marukilow model with default parameters in the gstacks module of Stacks 2.0.. This SNP panel was further filtered following additional criteria: *i*) genotyped in > 60% individuals; *ii*) minimum allele count (MAC) ≥ 3; *iii*) conformance to Hardy-Weinberg equilibrium (HWE), i.e. loci with significant (P < 0.05) F_IS_ values (Weir & Cockerham, 1984) detected in at least 25% of populations were removed; and *iv*) selection of the most polymorphic SNP in each RAD-tag.

### Genetic diversity and population structure

Genetic diversity (i.e. mean number of alleles per locus (Na), observed (Ho) and expected (He) heterozygosities), departure from Hardy–Weinberg equilibrium (HWE) and intrapopulation fixation index (F_IS_) were estimated for each bed using GENEPOP v4.0 (Rousset, 2008) and ARLEQUIN v3.5 (Excoffier & Lischer, 2010). Linkage disequilibrium between pairs of loci regarding physical distance was estimated with r^2^ across the cockle genome using the program Plink 1.9 (Chang et al., 2015; http://www.cog-genomics.org/plink/1.9) and its significance calculated through exact tests over genotypic contingency tables using GENEPOP v4.0.

Global and pairwise relative coefficients of population differentiation (F_ST_) between cockle beds were calculated with ARLEQUIN v3.5 using 10,000 permutations to test for significance. The number of genetically homogenous population units (K) was estimated using the variational Bayesian clustering method implemented in the package fastSTRUCTURE v2.3.4 (Raj et al., 2014) for K = 1 - 21, with an admixture ancestry model, convergence criterion of 1 × 10^−7^, five cross-validated sets and the simple prior (flat-beta prior). The most likely number of K was estimated using the “chooseK.py” program. This program gives the best K value and K corresponding with weak population structure in the data using the heuristic scores. Summarized outputs were carried out using the software DISTRUCT 1.1 (Rosenberg, 2004). Discriminant analyses of principal components (DAPC) were run in ADEGENET package (Jombart et al., 2010; Jombart & Ahmed, 2011) for the R platform (R Development Core Team, 2014; http://www.r-project.org). Data were transformed using PCA (Principal Component Analysis) and an appropriate number of principal components (PC) and discriminant functions (DF) were retained to explain > 90% of variance. Analyses of molecular variance (AMOVA) to study the distribution of genetic variation within (F_SC_) and among (F_CT_) bed groups obtained from fastSTRUCTURE were carried out with the program ARLEQUIN v.3.5 and their significance tested with 10,000 permutations. Isolation by distance (IBD) was evaluated through the correlation between geographical (measured as the shortest ocean distance) and genetic (measured as *F*_ST_/1-*F*_ST_; Rousset, 1997) distance matrices evaluated with a Mantel test with 10,000 permutations using NTSYS v.2.1 (Rohlf, 1993). All these analyses were performed both with the complete SNP dataset (9,309 markers) and the detected divergent outlier loci dataset (441 markers, see Results section).

### Outlier tests and gene mining

The Bayesian F_ST_-based method implemented in BAYESCAN v2.01 (Foll & Gaggiotti, 2008) was used to identify outlier loci potentially under selection in the whole Northeast Atlantic with samples grouped by locations. BAYESCAN was run using default parameters (i.e. 20 pilot runs; prior odds value of 10; 100,000 iterations; burn-in of 50,000 iterations and a sample size of 5,000). Loci with a False Discovery Rate (FDR, q-value) < 0.05 were considered outliers. RAD-tags including divergent outlier SNPs were mapped in the *C. edule* assembled genome (Tubío et al., unpublished) and used as landmarks for mining the genome to identify candidate genes related to selective drivers. To establish the genome windows for mining, linkage disequilibrium (LD) was evaluated across the whole genome and between consecutive outlier loci at specific genomic regions. It was expected that those regions under selection showed a higher LD than the average due to selective sweeping. LD was represented against physical distance for all pairs of markers across the whole genome, between markers separated up to 500 kb in each bed and up to 5,000 kb in the whole cockle collection of the Northeast Atlantic, respectively. The corresponding r^2^ values (the square of the correlation coefficient checking for the non-random association of alleles at pairs of loci) were averaged within 1 kb and 50 kb genomic windows within each bed and for the whole collection, respectively. Considering this information, and following a conservative criterion, we established windows of ± 200 kb around outlier markers in the selected genomic regions for mining (Supplementary Table 1). For this, the most consistent genomic regions related to divergent selection were selected according to the following criteria: *i*) the presence of two or more consecutive outliers in a region < 150 kb; and *ii*) the existence of a significant LD between them or above the maximum average genomic LD. Candidate genes for divergent selection included in these genomic windows were identified and annotated using the cockle transcriptome (Pardo et al., unpublished) and genome (Tubío et al., unpublished). Reactome and KEGG biological and molecular enriched pathways associated with these genes were retrieved from Kobas 2.0 (Xie et al., 2011) using *Homo sapiens* and *Drosophila melanogaster* as background because of their rich annotation and proximity, respectively.

### Landscape analyses

Genetic differentiation explained by the different spatial and abiotic factors was studied following a canonical redundancy analysis (RDA) using the VEGAN software (Oksanen, 2015) in R. For each bed (for those with temporal replicates, only information for the year 2017 was used), allele frequencies were estimated with ADEGENET package in R (Jombart & Ahmed, 2011) using the “makefreq” option applied on the ADEGENET “genpop” file. Loci with missing values were excluded from the analysis. Latitude and longitude together with the following abiotic factors were available for all the beds except for ASCE_17 (Sylt-Germany): sea surface temperature (SST, °C); sea bottom temperature (SBT, °C); sea surface salinity (SSS, psu); sea bottom salinity (SBS, psu); bottom shear stress (BSS, N·m^−2^); net primary productivity (NPP, mg·m^−3^·day^−1^) (Supplementary Table 2). Monthly information for all these abiotic factors was retrieved from the IBI_REANALYSIS_PHYS_005_002 model^1^ and the IBI_REANALYSIS_BIO_005_003 model^2^, for the period 2014-2018.

The nearest model cell with water was taken to extract the data. Then, both annual and spawning season (i.e. from April to August, see Malham et al., 2012) averages were calculated for each bed.

ANOVA was performed to test the significance of the variance associated to the different variables using 1,000 random permutations with VEGAN. Variance inflation factor (VIF) was estimated to explore collinearity (correlation) among landscape variables in the dataset. VIF values higher than 10 denote important collinearity problems, values from 5 to 10 moderate problems, while values lower than 5 indicate no collinearity problems (Marquardt, 1970). Different models were adjusted following a forward selection process with the PACKFOR package in R (Dray et al., 2009). This selection process corrects for highly inflated type I errors and overestimated amounts of explained variation (Vandamme et al., 2014). Thus, the reduced panel of explanatory variables is used to recalculate the total proportion of genetic variation in the variance partitioning. The weight of the different loci on the significant environmental vectors was obtained using VEGAN. All these analyses were performed separately for the complete and the divergent outlier SNP datasets.

### Larval dispersal modelling

A larval dispersal model was developed for the Northeast Atlantic in which virtual particles representing cockle larvae were ‘released’ from cockle bed locations along the Atlantic coast and transported by simulated ocean currents for the duration of their assigned pelagic larval phase. The particle trajectories were tracked enabling us to estimate the likely larval dispersal patterns and potential connectivity between different cockle populations.

Simulated ocean velocities were extracted from the IBI (Iberian Biscay Ireland) Ocean Analysis and Forecast system^3^. The underlying hydrodynamic model is based on the NEMO ocean model, version 3.6. The primitive equations are solved on a horizontal grid with a resolution of 1/36° and 50 unevenly spaced vertical levels with resolution decreasing from the surface into the deep. GEBCO08 was used for the bathymetry. Atmospheric forcing was extracted from ERA interim atmospheric fields. At the lateral open boundaries, ocean forcing fields were obtained from the global CMEMS GLOBAL eddy resolving system at 1/12°, and tidal forcing from the global FES2014 database (https://www.aviso.altimetry.fr/en/data/products/auxiliary-products/global-tide-fes.html). Freshwater fluxes from rivers were implemented for 33 rivers across the model domain. The model has been extensively validated and quality controlled (see Sotillo et al., 2015; 2020).

For the larval dispersal model, particles were released from 51 coastal sites (see Figure 1 for numbered release sites) covering 11 of the 14 wild natural beds sampled (11 because two sets of sample sites were in close proximity to one another, and one site was not within the ocean model domain), plus 40 other sites where cockles are known to habit. These 51 sites were roughly evenly distributed along the Atlantic coastline (ca. 60 - 150 km apart), hence reducing bias in the connectivity modelling from large differences in spatial distances between sites. From each site, cohorts of 400 particles were released on 01 April 2016. The cohorts of particles were transported by the simulated ocean currents using daily averaged current fields for an assigned pelagic larval phase of 40 days of which the last 10 days were used for analysis, thus accounting for uncertainties in the length of pelagic larval duration. The simulation was repeated at three different depth levels (1 m, 15 m and 30 m water depth); this strategy was designed to account for the large uncertainties regarding the vertical behaviour of *C. edule* larvae. This scenario was then repeated each day for the first 16 days in April to capture the variability in larval dispersal due to the tides. The procedure was then repeated for the first 16 days of months May to September, corresponding to the spawning phase of *C. edule* (Malham et al., 2012; Mahony et al., 2020), and then repeated for the years 2017 and 2018 to capture interannual variability from 2016 to 2018, due to changes in the formations, positions and strengths of the coastal and frontal currents. Therefore, a total of 17,625,600 ‘larvae’ particles were tracked for the experiment (400 particles × 51 sites × 3 depth levels × 16 release days × 6 months × 3 years).

Results are presented in terms of probability distribution maps and connectivity networks. To calculate connectivities, it was assumed that the larvae were able to settle during the last 10 days of their larval stage (i.e. days 31-40), therefore each particle’s positions during this time were used for the analysis of larval connectivities between sites. Larval connectivities were calculated as the percentage of larvae released from the source location arriving within a square of 0.2° latitude/longitude distance of one of the other 50 sink locations, or returning to the source location (self-recruitment). Connectivities were averaged over the three vertical release levels, but also presented separately in the Supplementary Materials.

## Results

### SNP genotyping and genetic diversity within beds

A total of ~2,000 M raw reads were initially analysed (~3.7 M reads per sample). After quality filtering and alignment, around 750 M reads were fed into Stacks to yield 315,744 loci with 726,911 SNP positions. Most of the SNPs filtered out were due to more than one match in the cockle’s genome. After filtering by population criteria, the number of retained SNPs for the whole population sample was 9,309. Nine individuals (two from Dee Estuary – WDE_17 and seven from Ria Formosa – PRF_18; Table 1) with low DNA quality and genotyping confidence (< 20% of SNPs genotyped) were removed. Thus, a total of 536 individuals from 14 beds were retained and used for subsequent analyses.

Genetic diversity was estimated as the average observed (Ho) and expected heterozygosity per locus (He). Ho ranged from 0.070 in Sylt – ASCE_17 and Burry – WBY_18 to 0.079 in Ría de Noia – SNO_17, Ria Formosa – PRF_17 and PRF_18 (mean ± SD = 0.074 ± 0.003), while He from 0.077 in Sylt – ASCE_17 and Burry – WBY_18 to 0.088 in SNO_17 (mean= 0.081 ± 0.003) (Table 1). F_IS_, which estimates the deviation from Hardy-Weinberg proportions within populations, was positive but not significant (i.e. heterozygote deficit; < 0.119; Table 1), so random mating sustains in all cockle populations. As expected, when a polymorphic criterion was considered (Minimum Allele Frequency (MAF) > 0.01) the number of polymorphic loci ranged from 1886 in Ria Formosa – PTE_18 to 4700 in Lombos do Ulla – SLO_17 (mean ± SD = 3455.9 ± 858.9) and accordingly, genetic diversity increased and its range enlarged: Ho from 0.134 in Lombos do Ulla – SLO_18 to 0.271 in Ria Formosa – PTE_18 (mean ± SD = 0.159 ± 0.030) and He from 0.153 in Lombos do Ulla – SLO_18 to 0.290 in Ria Formosa – PTE_18 (mean ± SD = 0.177 ± 0.030). This range widening was mainly due to the Tejo Estuary, Portugal, which represented an outlier population with both estimators much higher than in the other populations. Similar to the F_IS_ estimations considering all SNPs, the F_IS_ values using SNPs with the MAF > 0.01criterion were positive and non-significant in all cases (< 0.124; Table 1). Interestingly, the two Lombos do Ulla – SLO samples in Galicia severely affected by a *Marteilia cochilia* parasite outbreak showed the lowest genetic diversity (see Discussion).

### Outlier detection and gene mining

Detection of SNPs under selection was addressed with the BAYESCAN program, which tests whether the genetic differentiation between populations is above or below the neutral background (outlier loci), thus being suggestive of divergent or balancing selection, respectively. BAYESCAN analysis detected a total of 460 outlier loci potentially under selection, 19 under balancing selection and 441 under divergent selection in the whole Northeast Atlantic. The set of 441 loci under divergent selection was used along with the whole SNP dataset (9309 SNPs) for assessing patterns of structure under environmental drivers versus neutral factors (population size, larval dispersion) to identify candidate genes and functions associated with adaptation. All the outliers were mapped against the *C. edule* genome, mostly being distributed across the 19 mega-scaffolds of its assembly corresponding to the haploid chromosome number of the cockle’s karyotype (Insua & Thiriot-Quiévreux, 1992; Table 2). This information was used to estimate LD between adjacent markers regarding physical distance across the cockle’s genome within each population as well as in the whole cockle collection of the Northeast Atlantic (Fig. 2). Results showed that LD was low both within each population as well as in the whole studied area, and the highest average LD measured as r^2^ was always below 0.050, even for very short distances. Within each population LD was not significant for nearly all pairwise comparisons, in part due to the limited sample size (N ~ 30), but even when all data from the Northeast Atlantic was pooled (580 individuals), LD was mostly negligible and not significant above 50 kb on average. Using the whole population data, the most consistent genomic regions under divergent selection were defined by consecutive outlier markers at < 150 kb showing significant LD (P < 0.05) or above the maximum average in the whole studied area (r^2^ = 0.03; Fig. 2) and were located at mega-scaffold chromosome 1 (C1, four regions), C3 (two regions), and C2, C5, C10, C11 and C14 (one region) (Supplementary Table 1). These regions comprehended less than 25 kb and LD was significant in nearly all cases (P < 0.05), r^2^ averaging 0.166 and reaching up to 0.857 in one region of C1. These observations suggest selective sweeps at those regions, to say, selection of favourable haplotypes driven by particular environmental factors determining LD and/or loss of genetic diversity. Despite the poor annotation of the cockle’s genome outlined above, the windows evaluated in the selected genomic regions were particularly enriched in Gene Ontology (GO) terms related to neural function and development, immune response and defence, and metabolism and growth. Using the human genome as background, we could identify highly significant enriched molecular and metabolic pathways using Reactome and KEGG databases such as Immune System, Cellular Senescence and Necroptosis (corrected P-value < 0.001); Innate Immune System, Interleukin Signalling, Cell Cycle, Metabolism and Signal Transduction (P < 0.01), among other terms; and Insulin Secretion, Cytokine Signalling and Immune System and Lipolysis (P < 0.05), among others (Supplementary Table 3). Taking *Drosophila melanogaster* transcriptome as background, results were very poor due to the low number of significant homologies detected (not shown). Although information is still preliminary, results suggest essential biological functions related to adaptation in edible cockle that would deserve further work and that could be essential to understand the environmental factors driving selection in the Northeast Atlantic region.

**Table 2.** Distribution of outliers along the *Cerastoderma edule* genome assembly.

| Genomic Location | Bayescan Outliers (FDR < 0.05) |  |
| --- | --- | --- |
|  | Balancing selection | Divergent Selection |
| Mega-scaffold C1 | 2 | 40 |
| Mega-scaffold C2 | 1 | 33 |
| Mega-scaffold C3 |  | 39 |
| Mega-scaffold C4 | 1 | 31 |
| Mega-scaffold C5 | 2 | 28 |
| Mega-scaffold C6 |  | 24 |
| Mega-scaffold C7 | 2 | 13 |
| Mega-scaffold C8 | 3 | 24 |
| Mega-scaffold C9 | 2 | 18 |
| Mega-scaffold C10 | 1 | 23 |
| Mega-scaffold C11 | 1 | 15 |
| Mega-scaffold C12 |  | 22 |
| Mega-scaffold C13 | 1 | 25 |
| Mega-scaffold C14 | 2 | 14 |
| Mega-scaffold C15 | 1 | 16 |
| Mega-scaffold C16 |  | 32 |
| Mega-scaffold C17 |  | 13 |
| Mega-scaffold C18 |  | 4 |
| Mega-scaffold C19 |  | 11 |
| Other scaffolds (15) |  | 15 |
| Not found in genome |  | 1 |
| Total | 19 | 441 |

**Figure 2.**
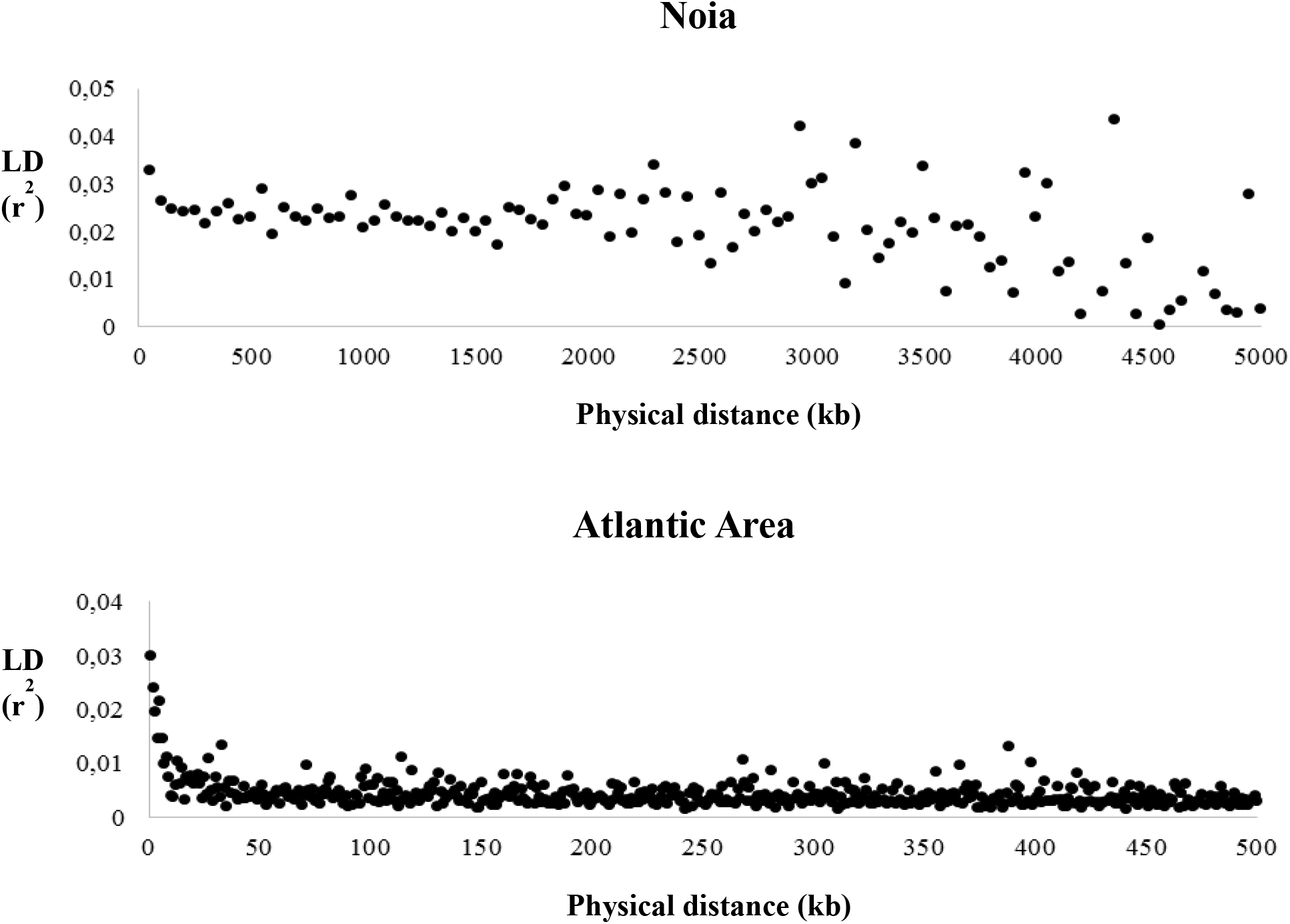
Linkage disequilibrium (r^2^) regarding physical distance between pairs of markers in the *Cerastoderma edule* genome in a representative cockle bed (Noia) and using the whole dataset from the Atlantic Area

### Population structure: temporal and geographical factors

All pairwise F_ST_ values between temporal replicates were non-significant (P > 0.050), suggesting temporal genetic stability between consecutive cockle’s cohorts in the Northeast Atlantic (Supplementary Table 4). This stability was confirmed when integrating the whole data set using an AMOVA analysis, where the percentage of variation associated to differences among temporal replicates within bed (F_SC_) was non-significant and negligible, while the percentage among sampling sites (F_CT_) was highly significant (P < 0.001) and higher than 3% (Model I, Table 3).

**Table 3.** AMOVA analyses for European *Cerastoderma edule* beds in the Atlantic Area.

|  | *F*-statistic | Variance component | % Variation |
| --- | --- | --- | --- |
| **Model I – Temporal (6 groups)** |  |  |  |
| Among beds ($F_{ST}$) | 0.03501*** | 358.724 | 3.50 |
| Among sampling sites ($F_{CT}$) | 0.03513*** | 359.951 | 3.51 |
| Among temporal replicates within sampling site ($F_{SC}$) | -0.00012 NS | -0.01227 | -0.01 |
| Within beds |  | 9.887.860 | 96.50 |
| **Model II – STRUCTURE - all dataset (K = 2)** |  |  |  |
| Among beds ($F_{ST}$) | 0.03769*** | 914.631 | 3.77 |
| Among fastSTRUCTURE groups ($F_{CT}$) | 0.02977*** | 722.427 | 2.98 |
| Among beds within fastSTRUCTURE groups ($F_{SC}$) | 0.00816*** | 192.204 | 0.79 |
| Within beds |  | 23.352.461 | 96.23 |
| **Model III – STRUCTURE - outliers (K =2 / K = 10)** |  |  |  |
| Among beds ($F_{ST}$) | 0.17195*** /<br>0.13055*** | 6.41246 /<br>4.63671 | 17.19/13.06 |
| Among fastSTRUCTURE groups ($F_{CT}$) | 0.13455*** /<br>0.12707*** | 5.01753 /<br>4.51316 | 13.45/12.71 |
| Among beds within fastSTRUCTURE groups ($F_{SC}$) | 0.04322*** /<br>0.00399*** | 1.39493 /<br>0.12355 | 3.74/0.35 |
| Within beds |  | 30.87961 /<br>30.87961 | 82.81/86.94 |
| NS: Non Significant ($P > 0.05$); ** $P < 0.01$; *** $P < 0.001$ | | | |

Pairwise F_ST_ values were significant between all the studied beds, except for some comparisons involving nearby sampled beds within each country (Supplementary Table 4). F_ST_ ranged from −0.0171 (Lombos do Ulla – SLO_17 vs Sado Estuary – PSA_19) to 0.0546 (Dee Estuary – WDE_17 vs Sado Estuary – PSA_19), with a global F_ST_ value of 0.0240 (P < 0.001). As expected, F_ST_ increased when only divergent outlier loci were considered (global F_ST_ = 0.1157, P < 0.001), ranging from −0.0133 (Dundalk Bay – IDC_18 vs Dundalk Bay – IDA_18) to 0.2216 (Dee Estuary – WDE_17 vs Sado Estuary – PSA_19). A consistent distribution of genetic diversity according to geographical distance was found, confirmed by a significant isolation by distance (IBD) pattern (complete dataset: *r* = 0.60541, P < 0.001; outlier dataset: *r* = 0.69568, P < 0.001; Supplementary Table 5).

Bayesian clustering analysis performed with fastSTRUCTURE using the complete dataset (i.e. 9,309 SNPs) rendered a value of K = 2 as the most probable population structure (Fig. 3). One group was formed by the northern beds (above 48° N, including the Bay of Somme (France) and north to Germany, Britain and Ireland), while the southern group was constituted by the beds near Arcachon (France) and south to Spain and Portugal (Fig. 3). AMOVA analysis using these two groups indicated that this structuring (F_CT_ = 2.98% of the total genetic diversity) captured close to the 80% of the total differentiation between populations (F_ST_ = 3.77%) (Model II, Table 3). The best K value with the outlier dataset was also 2 and the grouping was identical to that described with the complete dataset. However, the K value for the weak population structure using the heuristic scores provided by fastSTRUCTURE was 10. Among these, seven main groups were well defined: (*i*) North Sea and English Channel beds to the Bay of Somme (ASCE_17, NTX_18, FBS); (*ii*) the Dee bed in North Wales (WDE); (*iii*) Burry bed in South Wales (WBY); (*iv*) the Irish beds (IDA_18 and IDC_19); (*v*) the bed near Arcachon (FAR_17); (*vi*) the Spanish beds together with the northern Portuguese bed (SNO, SLO, PRA_17); (*vii*) the southern Portuguese beds (PTE_18, PSA_19, PRF) (Fig. 3).

**Figure 3.**
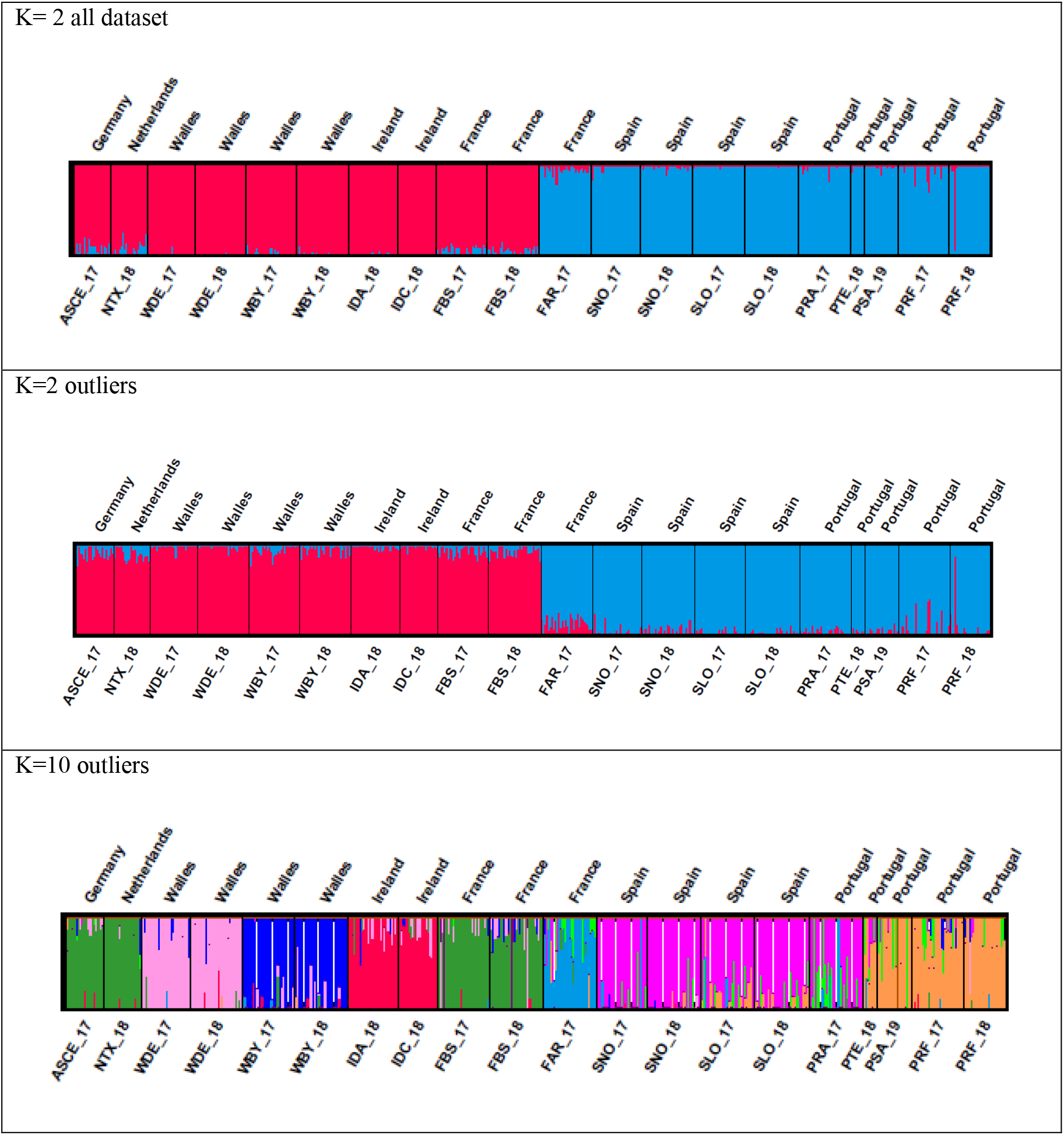
Population structure of *Cerastoderma edule* in the studied region using fastSTRUCTURE. Each vertical bar represents one individual, and the colour proportion for each bar represents the posterior probability of assignment of each individual to the different clusters (K) inferred by the program. Results obtained with K=2 using the complete dataset (A), K=2 using divergent outliers (B) and K=10 using divergent outliers (C) are represented. Codes are shown on Table 1.

AMOVA analyses with this outlier-based dataset (K = 10) assigned a higher percentage of genetic variation to differences among groups than with the whole dataset (Model III, Table 3). The percentage of variation associated to differences among beds within groups was the lowest (Table 3), confirming their genetic homogeneity, and accordingly, that the main differences among groups were captured (97.3% of the variation among beds). DAPC plots confirmed the results found with fastSTRUCTURE regarding the main north-south subdivision, but further clustering was suggested within groups. The analysis with the complete dataset showed an important dispersion within each group (Fig. 4A), while the analysis with the outlier dataset clearly identified four main differentiated groups: (*i*) Celtic and Irish Seas; (*ii*) North Sea and English Channel; (*iii*) the Bay of Biscay and Iberian waters to northern Portugal; and (*iv*) Iberian waters in southern Portugal, and even a more subtle subdivision up to the seven groups observed with STRUCTURE using outlier loci could be devised (Fig. 4B).

**Figure 4.**
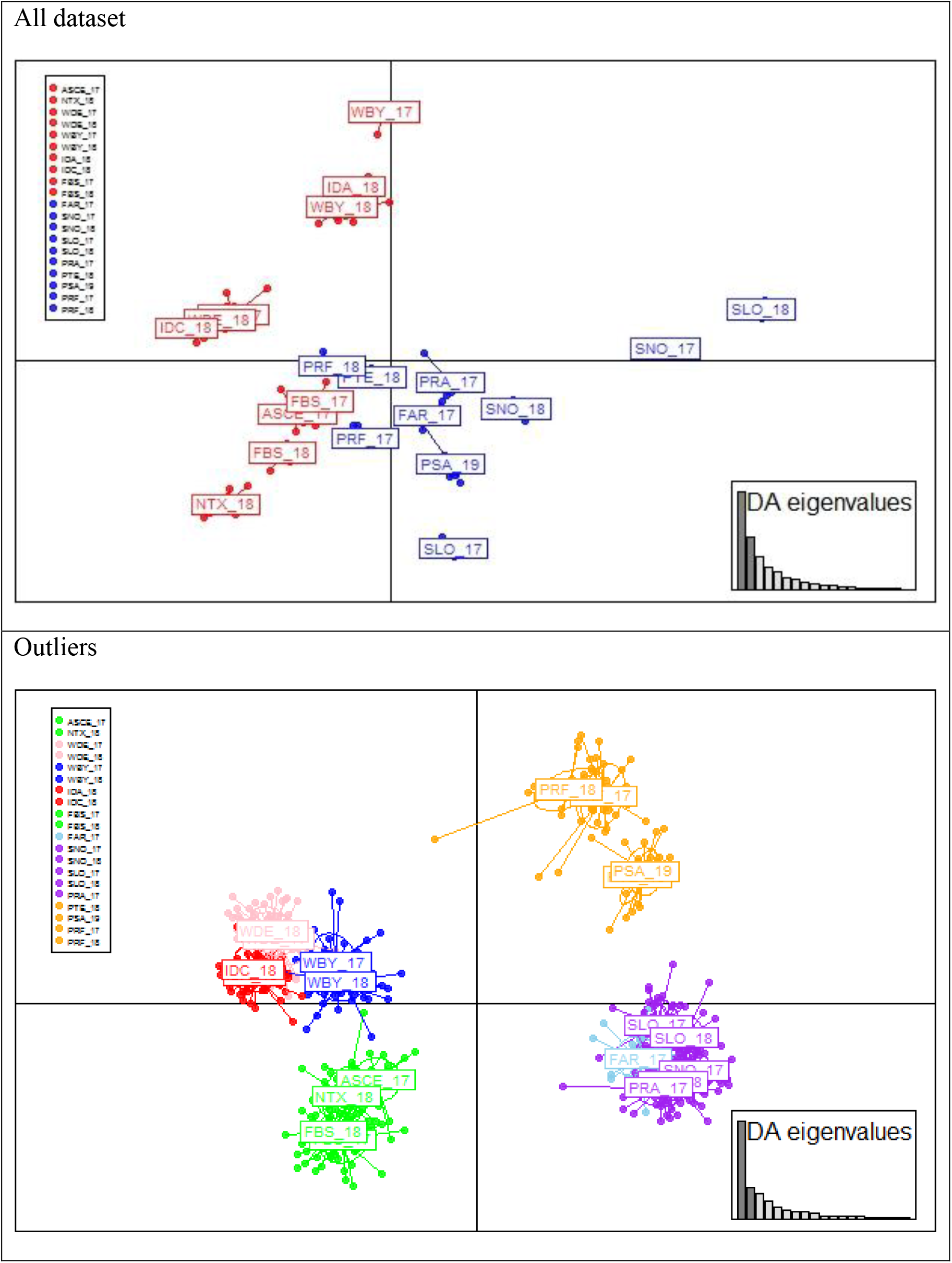
Discriminant Analysis of Principal Components (DAPC) plots of *Cerastoderma edule*. The weight of retained discriminant analysis (DA) eigenvalues to represent > 90% of variance are shown on right bottom box. Results using the complete dataset (A) and the divergent outliers (B) are represented. Codes are shown on Table 1

### Genetic-environmental associations

When all the environmental variables were included in the RDA analyses (measured as their average annual values for abiotic factors), only latitude was suggested as a driver for genetic differentiation, both for complete and outlier datasets (Table 4). Latitude was mainly associated with the first axis (Figs. 5A and 5C), which separated the beds in the two groups previously suggested by fastSTRUCTURE (i.e. northern and southern groups).

**Table 4.**
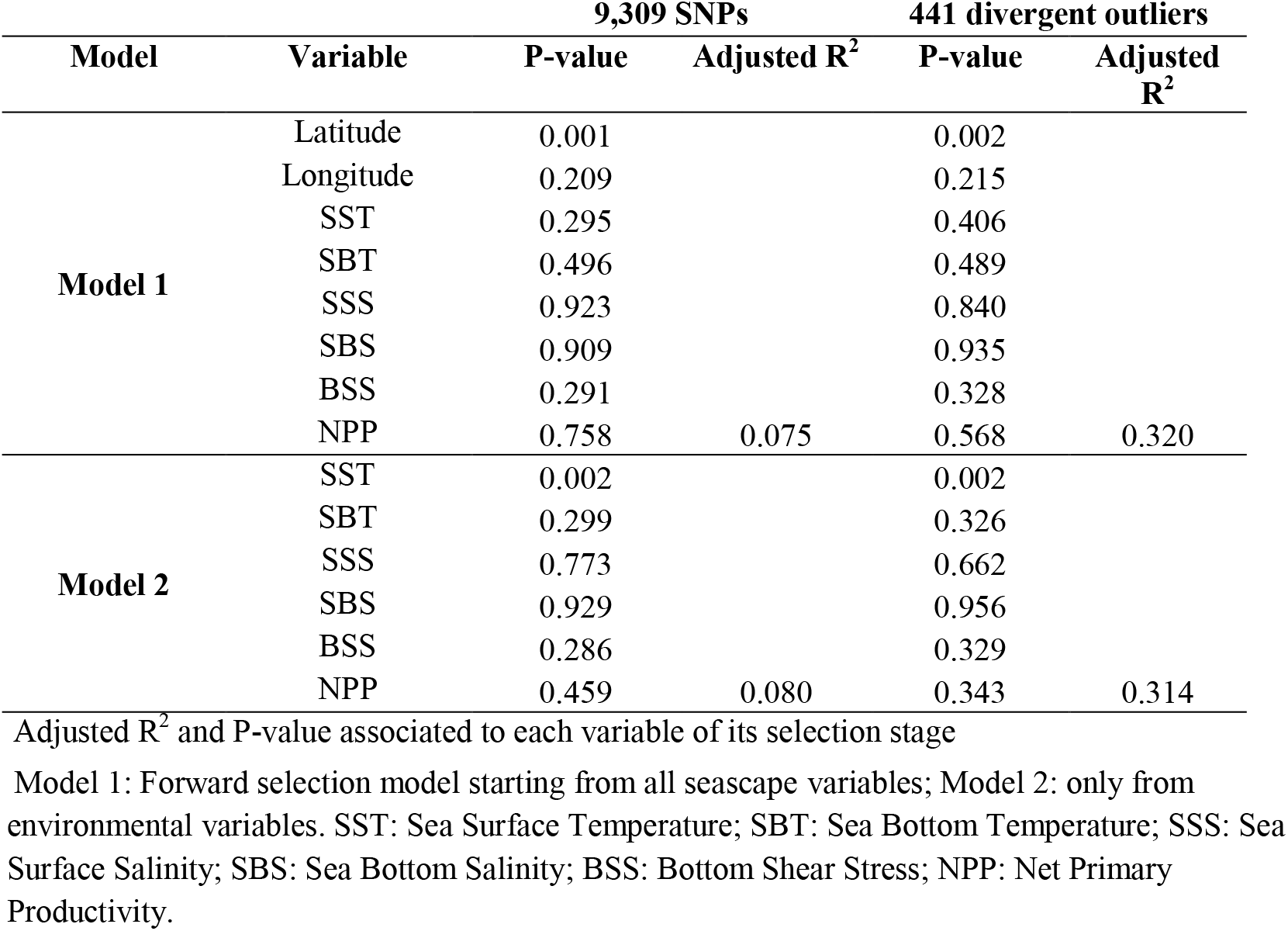
Results of the redundancy analysis (RDA) on *Cerastoderma edule* beds.

**Figure 5.**
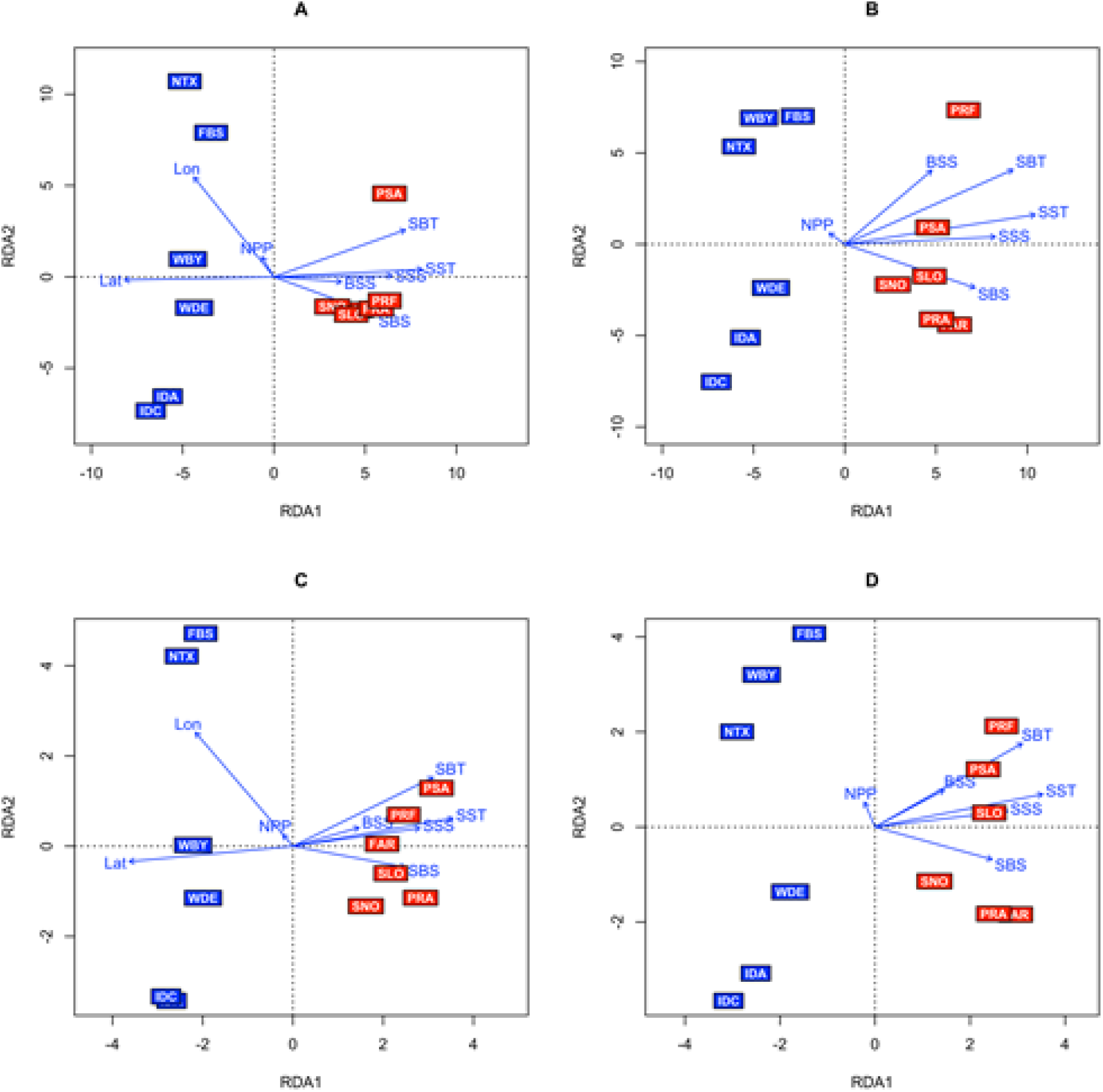
Redundancy analyses (RDA) of *Cerastoderma edule* samples from the studied area using the complete (A and B) and divergent outlier (C and D) datasets taking into account all landscape variables (Model 1, A and C) and only abiotic factors (Model 2, B and D). SST: Sea Surface Temperature; SBT: Sea Bottom Temperature; SSS: Sea Surface Salinity; SBS: Sea Bottom Salinity; BSS: Bottom Shear Stress; NPP: Net Primary Productivity.

Despite being non-significant, the remaining variables were also associated with the first axis excluding longitude and net primary productivity (NPP), which were associated with the second axis. When exclusively abiotic variables were used, the sea surface temperature (SST) was suggested as a significant driver (Table 4). SST was also related to the first axis and again separated the cockle beds in the northern and southern groups (Fig. 5B and 5D). However, these results should be taken with caution since VIF values associated to most variables were > 20 excluding longitude (VIF = 0.55), bottom shear stress (BSS; VIF = 4.70) and net primary production (NPP; VIF = 5.43), so we decided to evaluate them individually with longitude, BSS and NPP. All those variables were significant with both SNP datasets (P < 0.05), excluding bottom temperature (SBT), and related to differentiation between the northern and southern groups (Supplementary Table 6). Therefore, several environmental variables would be likely shaping the cockle genome in the Northeast Atlantic, including salinity (SSS and SBS), but all of them appeared highly correlated splitting the studied area into northern and southern groups, as observed with the whole SNP dataset. The results obtained using only the spawning season values for the abiotic factors were identical to those found with the average annual values (data not shown).

### Current modelling and larval dispersal

Connectivity pathways from the larval dispersal modelling, time-averaged and averaged across all three depth scenarios, are shown in Figure 6. All sites along the coasts of Portugal, Spain and the west coast of France up to Brittany (sites 1 – 19) were simulated to be potentially well connected to multiple neighbouring sites along the coastline with relatively high levels of connectivity (>1%). Even if simulated connectivity between neighbouring sites was less than 1%, generally multiple connection pathways were simulated. Across the headland of Brittany into the English Channel, where genetic differentiation was established, the modelling simulated weak connectivities (less than 0.2% between sites 18/19 and site 20, on average, with several scenarios simulating no connectivity). Sites within the English Channel were simulated to be well connected (especially the eastern sites) with multiple connectivity pathways between sites on either side of the channel and into the North Sea. Similar to Brittany, we simulated relatively weak (and one-way only) connectivity from the English Channel to the Celtic Sea, around the headland of SW Britain (connecting site 33 to 34). Sites within the Celtic Sea were simulated to be well connected: greater than 1% between many sites and with notable larval transport along the Celtic Sea Front potentially connecting English/Welsh sites with Ireland. Along the south and west coast of Ireland, particles became entrained in the Irish Coastal Current and were transported westwards then northwards along the coast, thus simulating potentially strong connectivities between sites 41 to 51.

**Figure 6.**
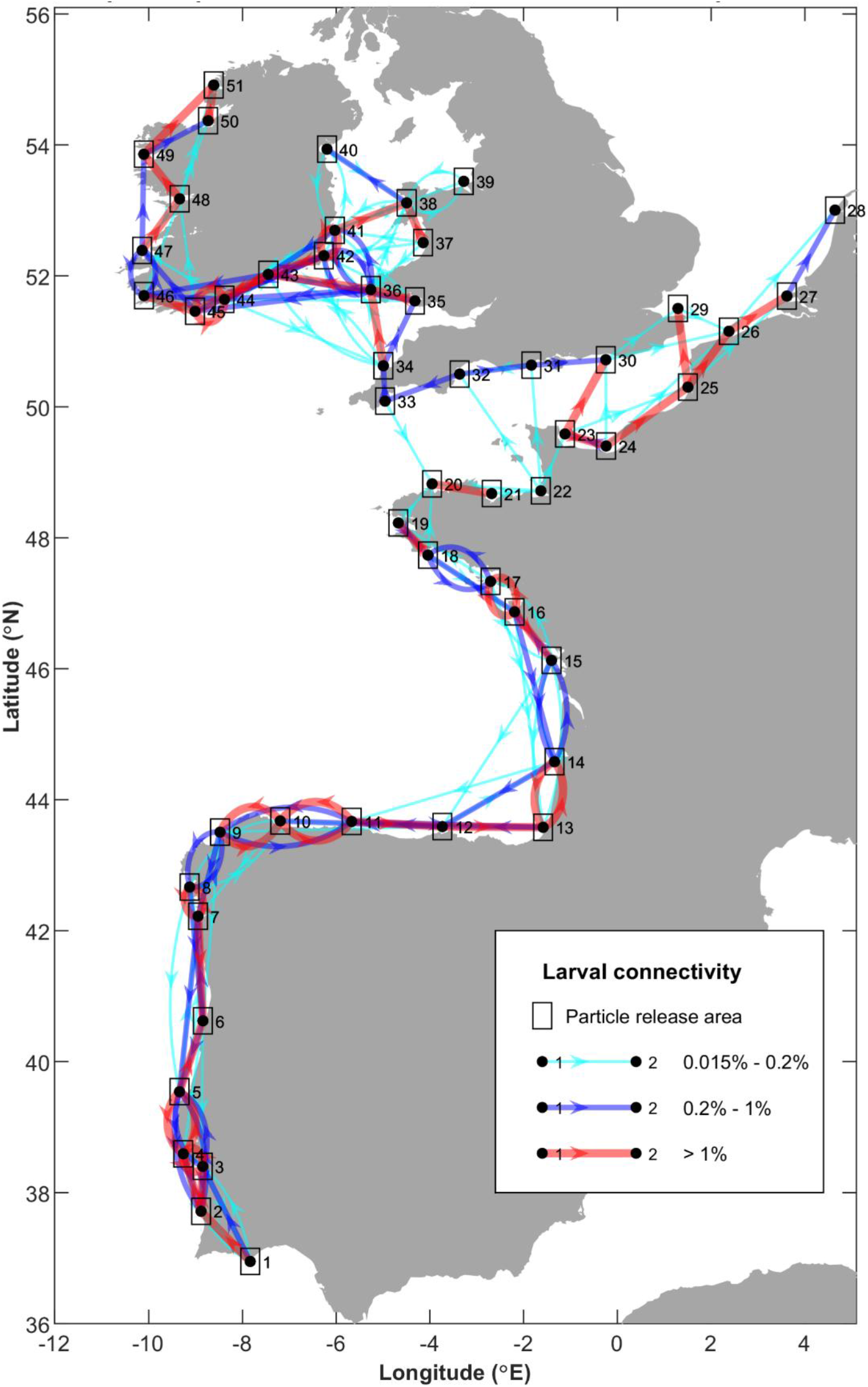
Mean larval connectivity pathways for April to September from 2016 – 2018 releases at depths of 1, 15 and 30 m. The direction of the arrows indicates the direction of larval transport and the color and thickness of the connection display the strength of the connection.

Taken individually, the larval behaviour scenarios (i.e. particle cohorts released (and fixed) at 1m, 15m and 30m water depths) showed considerable differences in the simulated connectivity networks. The surface (1m) release scenario (Supplementary Fig. 1) simulated the highest degree of connectivity with almost all sites receiving larvae from at least two other nearby sites and sites being interlinked along most coast lines. The 15m scenario (Supplementary Fig. 2) simulated a similar picture, with a well-connected Portuguese and Bay of Biscay coastline in particular, and a well-connected Irish and Celtic Sea region. However, simulated connections from the French Atlantic coast into the English Channel and from the English Channel into the Celtic Sea were much weaker (less than 0.2%, on average, with periods of no connectivity). For the 30m scenario (Supplementary Fig. 3), we simulated the same areas to be well-connected, but no dispersal was simulated between southern and northern Brittany, nor between southern and northern Cornwall, thus isolating the Atlantic sites, the English Channel and the Celtic/Irish Sea sites from each other. The simulated interannual variations in connectivities were greatest for the surface release scenario and decreased the deeper the larvae were in the water column (results not shown). For the 15m and 30m depth scenarios, simulated connectives over different years tended to vary in strength but much less so in overall patterns. Overall, the larval dispersal modelling suggested that the sites may be grouped into three distinct cohorts: (1) the Portuguese, Spanish and French Atlantic coast sites; (2) the English Channel sites; and (3) and the Celtic and Irish Sea sites.

## Discussion

### Genetic diversity

Maintenance of genetic diversity is crucial for the adaptation of natural populations to environmental changes. Preservation of connectivity among populations is essential to maintain the genetic diversity within and between populations to counteract the threat of demographic depletion caused by overexploitation or diseases (Frankham et al., 2002). Moreover, genetic diversity existing in the wild is important for commercial species, such as *C. edule*, to support breeding programs for selecting high-value seed for intensive production in particular areas (Lallias et al., 2010; do Prado et al., 2018; Petereit et al., 2018; Vera et al., 2019). Genetic diversity was similar across all cockle beds studied in the Northeast Atlantic, especially when no polymorphic filtering criterion was applied (MAF; He range: 0.077 −0.086). When a MAF > 0.01 filtering was applied to avoid the bias of low polymorphic loci in accordance with the sample size managed (i.e. 1/(2*highest N) = 0.016; N (SLO_18) = 32 individuals on average), the genetic diversity range widened (0.153 - 0.290), but mostly due to the Tejo Estuary representing an outlier population with much higher He than the rest (He = 0.290). There is not a straightforward explanation for the high genetic diversity in Tejo Estuary, such as admixture of differentiated stocks in that area (Wahlund effect; see below), since the intrapopulation fixation index F_IS_ was not significant and even among the lowest in the samples studied. At the other end of the range, Lombos do Ulla (Galicia) samples showed the lowest genetic diversity in all the Northeast Atlantic, but in this case the serious population depletion caused by the parasite *Marteilia cochillia* that affects this bed since 2012 (Villalba et al., 2014), could be responsible. Moreover, all populations showed a consistent slight heterozygote deficit (F_IS_ > 0), although non significant, likely related to a technical genotyping issue associated with the presence of null alleles as previously reported in molluscs (Pino-Querido et al., 2014).

Heterozygosity figures in our study were similar to those reported by Coscia et al., 2020 (He: 0.144 - 0.156), who also applied a MAF > 0.01 filter to characterise genetic diversity in *C. edule* beds from the Celtic and Irish Seas (i.e. southeast Ireland and Wales) using a classical RADseq technique (Etter et al., 2011). Genetic diversity observed for *C. edule* in our study was similar to that reported for the sister species *C. glaucum* (He: 0.078 - 0.137; Sromek et al., 2019); however, both cockle species showed lower genetic diversity at genomic scale than other bivalves such as *Placopecten magellanicus* (He = 0.271 ± 0.133, Van Wyngaarden et al., 2017), *Crassostrea virginica* (He ~ 0.300; Bernatchez et al., 2019), *Ostrea edulis* (He ~ 0.300; Vera et al., 2019) and *Crassostrea gigas* (He ~ 0.300; Gutierrez et al., 2017), suggesting the lower genetic diversity to be a particular feature of the genus *Cerastoderma*, although we cannot discard that the different methodologies applied for SNP genotyping in those studies might explain this observation.

Genetic diversity (He) showed slight but significant differences between the two major northern and southern cockle genetic groups identified in the Northeast Atlantic (N vs S = − 0.0032; Mann-Whitney U tests; P = 0.031), but not when the MAF > 0.01 filtering was applied (P > 0.05). Higher genetic diversity would be expected in lower latitude populations of the Northern Hemisphere because of their role as glacial refugia during Quaternary glaciation (Hewitt, 2000). Previous studies with microsatellite loci did not detect differences between both groups (Martínez et al., 2013), and even supported higher diversity in the Celtic/Irish Seas, English Channel and the North Sea (Martínez et al., 2015) corresponding to the present northern group. Moreover, higher genetic diversity has been found in northern beds from the Fennoscandian region and Russia, using mtDNA markers, which suggested a cryptic northern glacial refugia for *C. edule* (Krakau et al., 2012, Martínez et al., 2015), which has also been described for other marine species (Luttikhuizen et al., 2008; Maggs et al., 2008; Sotelo et al., 2020). Our extensive genome-wide analysis indicates slight differences in genetic diversity for the edible cockle in the whole Northeast Atlantic evaluated, suggesting notable connectivity among samples.

### Population structure and connectivity

Knowledge of population structure, including local adaptations, is crucial to define and apply sustainable strategies for the management and conservation of exploited species (Frankham et al., 2002; Bernatchez et al., 2017). For *C. edule,* the presence of at least two main population units within its natural range, delimited by the western English Channel, had been reported in previous studies (Krakau et al., 2012; Martínez et al., 2015) and here has been confirmed with a more in depth population genomic analysis using the whole SNP dataset. The northern group in our study, comprising Celtic/Irish Sea, English Channel and North Sea beds, corresponds with the northern group defined by Martínez et al., (2015). This structure has also been documented in other mollusc species with a similar distribution range such as *O. edulis* (Vera et al., 2016). The results from our larval dispersal modelling, based on potential ocean-driven larval flows between sites along the Northeast Atlantic, also suggests a major discontinuity at the French Brittany headland, indicating that the oceanography of the Northeast Atlantic plays an important role in shaping the genetic structure of cockles. The frontal systems (Ushant Front and Celtic Sea Front) when fully established seem to represent barriers to dispersal, thus limiting e.g. the transport of larvae into the English Channel and into the Irish Sea. Headlands, such as Brittany and Cornwall, also appear to act as barriers, potentially due to diverging current systems.

Genomic scans aid detecting footprints of selection regarding the neutral background, which represents invaluable information for sustainable management and conservation of fisheries (Nielsen et al., 2009; Bernatchez, 2016). Genetic markers showing a significant departure, above or below the neutral dataset are potential outliers under divergent or stabilizing selection, respectively, which can unravel a fine-scale structure related to environmental variables critical for resources management (Vera et al., 2019; Coscia et al., 2020). Temperature, salinity and other abiotic factors have been suggested as potential drivers for adaptive differentiation in *C. edule* (Coscia et al., 2020), as in other marine species in the region (Vera et al., 2016; do Prado et al., 2018). In our study, the RDA analyses confirmed the effect of temperature in a wider geographical range, the major genetic subdivision north-south in the Northeast Atlantic being associated with a latitudinal annual mean temperature gradient of ~5.5 °C between the warmest (Ria Formosa) and the coldest station (Dundalk).

Moreover, the analysis with outlier loci enabled us to disclose a significant substructure in the Northeast Atlantic beyond the two major groups identified with the whole dataset. Thus, the northern Atlantic group appeared also split into two major subgroups including the Celtic/Irish Sea beds in the west and the English Channel and North Sea beds in the east using outlier information. Concurrently, this northern subdivision was also detected with the larval dispersal modelling performed here. Within the southern group, Portuguese samples were subdivided in a southern group (Ria Formosa – PRF, Tejo Estuary – PTE_18, Sado Estuary – PSA_19) differentiated from the northern sample (Aveiro – PRA_17), which clustered with northwest Spanish beds (SNO, SLO). In the larval dispersal modelling, very weak connectivity was also observed between both Portuguese subgroups during June and July at depths of 15 m and 30 m (see Supplementary Figs. 4 and 5), but more data on the spawning period are needed to confirm the role of ocean currents on the observed genetic subdivision (see e.g. Mahony et al., 2020). In fact, strong connectivities were detected during the other simulated months. Northwards, Cape Finisterre has been suggested as a biogeographical barrier for marine organisms (López-Jamar et al., 1992; Abaunza et al., 2008; Piñeira et al., 2008) including *C. edule*, separating northwest Spanish beds from those in the Bay of Biscay (Martínez et al., 2013). Unfortunately, the Spanish beds in the Bay of Biscay were not analysed, but the Arcachon bed (FAR_17) in that Bay was significantly differentiated from the northwest Spanish group, suggesting a possible barrier between the Galician and Southern France beds, potentially Cape Finisterre. However, this substructure did not appear to be related to dispersal limitations of larvae according to our modelling data and more detailed sampling should be carried out in Biscay Bay to confirm this hypothesis. In fact, Cape Finisterre does not appear to represent a barrier to gene flow in other species (Domingues et al., 2010; López et al., 2015; Riquet et al., 2019).

As outlined above, the genetic structure of edible cockle in the Northeast Atlantic shows a notable correspondence with the larval dispersal modelling performed here, although some discordances are suggested depending on the set of markers used (neutral vs outlier loci). However, both datasets might be fully compatible if other variables such as larval depth in the water column is considered in larval dispersion modelling. The simulations carried out with larvae located at three different depths suggest that deeper depths (15 m and 30 m) correlate better with the major subdivision observed with neutral loci (north vs. south) and are in accordance with other studies on bivalve larval behaviour which show that larvae are preferably located below the surface of the water column rather than close to it (e.g. Scrope-Howe & Jones, 1986; Irigoen et al., 2004). This result also suggests the importance of oceanographic studies of larval distributions throughout the water column, and that knowledge on larval behaviour cannot be solely gained from tank experiments under laboratory conditions. Large uncertainties still persist since the larval behaviour of *C. edule* with regard to swimming behaviour remains to be studied in detail. Whilst the surface releases showed a higher degree of interannual variability, the deeper releases had stable connectivity patterns between years, thus tying in with the genetic structure which seems stable interannually.

A final thought to understand larval dispersion and environmental drivers shaping the cockle’s genome is why both factors split similarly the Northeast Atlantic area when they should have a distinct influence on genetic diversity. It would be expected that ocean current patterns and fronts were more related to neutral genetic variation, while outlier loci were with environmental factors driving selection. However, the difficulty to disentangle the effects of neutral and outlier loci datasets when defining the major subgroups of cockles, suggests a correlation between dispersal patterns and environmental variables. This is clear when considering the two groups defined by the major barrier at French Brittany and the north-south variation in temperature, but it is likely that ocean current patterns and seasonally-generated frontal flows may too be structuring environmental ecosystems and hence the correlation observed, not only at macrogeographic level, but also at a meso-scale. A refined analysis of genetic diversity at a local scale around the main fronts and physical barriers described here would provide essential information to understand the observed correlations.

### Fisheries management

Our results provide useful information for the management of cockle beds in the Northeast Atlantic and could be valuable for obtaining suitable seed either for restocking of depleted populations or for finding broodstock to enhance cockle production. An improved definition of management units considering both demography and adaptation to environmental variation along the Northeast Atlantic can now be delineated, allowing the future definition of adaptive management units (AMU, Bernatchez et al., 2017). Two main operational units, located northwards and southwards French Brittany can be defined as the basic proposal for management, but a more ambitious approach should include at least five different units: (*i*) Irish/Celtic Sea; (*ii*) English Channel/North Sea; (*iii*) Bay of Biscay; (*iv*) northwest Spain and north Portugal; and (*v*) south Portugal. Further, our data and those from Coscia et al. (2020) suggest that the Irish/Celtic Sea unit could be additionally refined. The information from this study might be useful to define sets of markers, starting from outlier loci, which could be applied to found brookstock for restocking depleted populations and to track individuals to their units that could aid the identification of illegal transferences between countries or from disease-affected areas. Our data represents the baseline to monitor restocking and to evaluate the impact of intensive aquaculture on cockle beds. Further work should be done at a more refined scale combining genetic and larval advection patterns around the main areas of differentiation in the Northeast Atlantic suggested in this study.

## Acknowledgements

The research leading to these results has received funding from the Interreg Atlantic Area Programme through the European Regional Development Fund for the project Co-Operation for Restoring CocKle SheLlfisheries and its Ecosystem Services in the Atlantic Area (COCKLES, EAPA_458/2016), www.cockles-project.eu. Authors wish to thank L. Insua, S. Sánchez-Darriba and S. Gómez for their technical support. Authors are also indebted with COCKLES, VIVALDI and Scuba Cancers partners, together with M.L. Conde-Varela, M.T. Fernández-Núñez, M. García-Graña, P. Luttikhuizen, S. Pereira, A. Simón, and L. Solís for providing many of the samples analyzed. Supercoputing Centers of Galicia (www.cesga.es) and Wales (www.supercomputing.wales) provided computing facilities and technical support for the genotyping and larval dispersal modelling, respectively.

## Footnotes

1 https://resources.marine.copernicus.eu/?option=com_cswandtask=results?option=com_cswandview=detailsandproduct_id=IBI_REANALYSIS_PHYS_005_002

2 https://resources.marine.copernicus.eu/?option=com_cswandtask=results?option=com_cswandview=detailsandproduct_id=IBI_REANALYSIS_BIO_005_003

3 IBI_ANALYSIS_FORECAST_PHYS_005_001; see https://resources.marine.copernicus.eu/?option=com_cswandview=detailsandproduct_id=IBI_ANALYSIS_FORECAST_PHYS_005_001 for details and to download the data.

## Notes

### Competing Interest Statement

The authors have declared no competing interest.

## References

Abaunza, P., Murta, A. G., Campbell, N., Cimmaruta, R., Comesana, A.S., Dahle, G García Santamaría, M.T., Gordo, L.S., Iversen, SA., MacKenzie, K., Magoulas, A., Mattiucci, S., Molloy, J., Nascetti, G., Pinto, A.L., Quinta, R., Ramos, P., Sanjuan, A., Santos, A.T., Stransky, C., Zimmermann, C. (2008). Stock identity of horse mackerel (*Trachurus trachurus*) in the Northeast Atlantic and Mediterranean Sea: Integrating the results from different stock identification approaches. Fisheries Research, 89, 196–209. doi:10.1016/j.fishres.2007.09.022

Beaumont, A.R., Day, T.R., Gade, G. (1980). Genetic variation at the octopine dehydrogenase locus in the adductor muscle of *Cerastoderma edule* (L.) and six other bivalve species. Marine Biology Letters, 1, 137–148.

Bernatchez, L. (2016). On the maintenance of genetic variation and adaptation to environmental change: Considerations from population genomics in fishes. Journal of Fish Biology, 89, 2519–2556. doi.org/10.1111/jfb.13145

Bernatchez, L., Wellenreuther, M., Araneda, C., Ashton, D.T., Barth, J.M.I., Beacham, T.D., Maes, G.E., Martinsohn, J.T., Miller, K.M., Naish, K.A., Ovenden, J.R., Primmer, C.R., Suk, H.Y., Therkildsen, N.O., Withler, R.E. (2017). Harnessing the power of genomics to secure the future of seafood. Trends in Ecology & Evolution, 32, 665–680. doi:10.1016/j.tree.2017.06.010

Bernatchez, S., Xuereb, A., Laporte, M., Benestan, L., Steeves, R., Laflamme, M., Bernatchez, L., Mallet, M.A. (2019). Seascape genomics of eastern oyster (*Crassostrea virginica*) along the Atlantic coast of Canada. Evolutionary Applications, 12, 587–609. doi:10.1111/eva.12741.

Blanco-González, E., Knutsen, H., Jorde, P.E. (2016). Habitat discontinuities separate genetically divergent populations of a rocky shore marine fish. Plos One, 11, e0163052. doi.org/10.1371/journal.pone.0163052

Brown, J., Carrillo, L., Fernand, L., Horsburgh, K. J., Hill, A.E., Young, E.F., Medler, K.J. (2003). Observations of the physical structure and seasonal jet-like circulation of the Celtic Sea and St. George’s Channel of the Irish Sea. Continental Shelf Research, 23, 533–561. doi:10.1016/s0278-4343(03)00008-6

Burgess, S.D., Bowring, S., Shen, S.Z. (2014). High-precision timeline for Earth’s most severe extinction. Proceedings of the National Academy of Sciences USA, 111, 3316–3321. doi.org/10.1073/pnas.1317692111

Catchen, J., Hohenlohe, P.A., Bassham, S., Amores, A., Cresko, W.A. (2013). Stacks: an analysis tool set for population genomics. Molecular Ecology, 22, 3124–3140. doi:10.1111/mec.12354

Chang, C.C., Chow, C.C. Tellier L., Vattikuti, S., Purcell, S.M., Lee, J.J. (2015) Second-generation PLINK: rising to the challenge of larger and richer datasets GigaScience, 4, 7. doi:10.1186/s13742-015-0047-8

Charria, G., Lazure, P., Le Cann, B., Serpette, A., Reverdin, G., Louazel, S., Batifoulier, F., Dumas, F., Pichon, A., Morel, Y. (2013). Surface layer circulation derived from Lagrangian drifters in the Bay of Biscay. Journal of Marine Systems, 109, S60–S76. doi.org/10.1016/j.jmarsys.2011.09.015

Clucas, G. V., Lou, R. N., Therkildsen, N. O., Kovach, A. I. (2019). Novel signals of adaptive genetic variation in northwestern Atlantic cod revealed by whole-genome sequencing. Evolutionary Applications, 12, 1971–1987. doi.org/10.1111/eva.12861

Coscia, I., Wilmes, S.B., Ironside, J.E., Goward-Brown, A., O’Dea, E., Malham, S.K., McDevitt, A.D., Robins, P.E. (2020). Fine-scale seascape genomics of an exploited marine species, the common cockle *Cerastoderma edule*, using a multimodelling approach. Evolutionary Applications, 13, 1854–1867. doi:10.1111/eva.12932

Cowen, R.K., Sponaugle, S. (2009). Larval dispersal and marine population connectivity. Annual Review of Marine Science, 1, 443–466. doi.org/10.1146/annurev.marine.010908.163757

Danancher, D., Garcia-Vazquez, E. (2011). Genetic population structure in flatfishes and potential impact of aquaculture and stock enhancement on wild populations in Europe. Reviews in Fish Biology and Fisheries, 21, 441–462. doi.org/10.1007/s11160-011-9198-6

Dare, P.J., Bell, M.C., Walker, P., Bannister, R.C.A. (2004). Historical and current status of cockle and mussel stocks in The Wash. CEFAS, Lowestoft

de Montaudouin, X., Bachelet, G., Sauriau, P.G. (2003). Secondary settlement of cockles Cerastoderma edule as a function of current velocity and substratum: a flume study with benthic juveniles. Hydrobiologia, 503, 103–116. doi:10.1023/B:HYDR.0000008493.83270.2d

do Prado, F.D., Vera, M., Hermida, M., Bouza, C., Pardo, B.G., Vilas, R., Blanco, A., Fernández, C., Maroso, F., Maes, G., Turan, C., Volckaert, F., Taggart J., Carr, A., Ogden, R., Nielsen, E.E., the Aquatrace consortium, Martínez P. (2018). Parallel evolution and adaptation to environmental factors in a marine flatfish: Implications for fisheries and aquaculture management of the turbot (*Scophthalmus maximus*). Evolutionary Applications, 11, 1322–1341. doi:10.1111/eva.12628

Domingues, C.P., Creer, S., Taylor, M.I., Queiroga, H., Carvalho, G.R. (2010) Genetic structure of *Carcinus maenas* within its native range: larval dispersal and oceanographic variability. Marine Ecology Progress Series, 410, 111–123. doi.org/10.3354/meps08610.

Dray, S., Legendre, P., & Blanchet, G. (2009). “PACKFOR: Forward Selection with permutation (Canoco p. 46).” R package version 0.0-7/r58 (2009).

Etter, P., Bassham, P., Hohenlohe, P., Johnson, E., Cresko, W. (2011). SNP discovery and genotyping for evolutionary genetics using RAD sequencing. Methods in Molecular Biology, 772, 157–178. doi:10.1007/978-1-61779-228-1_9

Excoffier, L., Lischer, H.E.L. (2010). Arlequin suite ver 3.5: a new series of programs to perform population genetics analyses under Linux and Windows. Molecular Ecology Resources, 10, 564–567. doi:10.1111/j.1755-0998.2010.02847.x

Fernand, L., Nolan, G.D., Raine, R., Chambers, C.E., Dye, S.R., White, M., Brown, J. (2006). The Irish coastal current: A seasonal jet-like circulation. Continental Shelf Research, 26, 1775–1793.

Foll, M., Gaggiotti, O. (2008). A genome-scan method to identify selected loci appropriate for both dominant and codominant markers: A Bayesian perspective. Genetics, 180, 977–993. doi:10.1534/genetics.108.092221

Frankham, R., Ballou, J.D., Briscoe, D.A. (2002). Introduction to conservation genetics. Cambridge: Cambridge University Press.

Galparsoro, I., Borja, A., Uyarra, M.C. (2014). Mapping ecosystem services provided by benthic habitats in the European North Atlantic Ocean. Frontiers in Marine Science, 1, 23. doi:10.3389/fmars.2014.00023

Gutierrez, A.P., Turner, F., Gharbi, K., Talbot, R., Lowe, N.R., Peñaloza, C., McCullough, M., Prodöhl, P.A., Bean, T.P., Houston, R.D. (2017). Development of a Medium density combined-species SNP array for Pacific and European oysters (*Crassostrea gigas* and *Ostrea edulis*). G3-Genes Genomes Genetics, 7, 2209–2218. doi:10.1534/g3.117.041780

Hayward, P. J., & Ryland, J. S. (1995). Handbook of the marine fauna of north-west Europe. Oxford University Press, Oxford

Hewitt, G. (2000). The genetic legacy of the Quaternary ice ages. Nature, 405, 907–913. doi.org/10.1038/35016000

Honkoop, P.J.C., van der Meer, J. (1998). Experimentally induced effects of water temperature and immersion time on reproductive output of bivalves in the Wadden Sea. Journal of Experimental Marine Biology and Ecology, 220, 227–246. doi:10.1016/s0022-0981(97)00107-x

Horsburgh, K.J., Hill, A.E., Brown, J. (1998). A summer jet in the StGeorge’s Channel of the Irish Sea. Estuarine Coastal and Shelf Science, 47, 285–294. doi:10.1006/ecss.1998.0354

Hughes, A. R., Hanley, T. C., Byers, J. E., Grabowski, J. H., McCrudden, T., Piehler, M. F., Kimbro, D. L. (2019). Genetic diversity and phenotypic variation within hatchery-produced oyster cohorts predict size and success in the field. Ecological Applications, 29, e01940. doi:10.1002/eap.1940

Hummel, H., Wolowicz, M., Bogaards, R.H. (1994). Genetic variability and relationships for populations of *Cerastoderma edule* and of *C*. *glaucum*complex. Netherlands Journal of Sea Research, 33, 81–89. doi:10.1016/0077-7579(94)90053-1

Insua, A., Thiriot-Quiévreux, C. (1992). Karyotypes of *Ceratoderma edule*, *Venerupis pullastra* and *Venerupis rhomboides* (Bivalvia, Veneroida). Aquatic Living Resources, 5, 1–8. doi.org/10.1051/alr:1992001

Irigoien, X., Conway, D.V., Harris, R.P. (2004). Flexible diel vertical migration behaviour of zooplankton in the Irish Sea. Marine Ecology Progress Series, 267, 85–97.

Jiménez-Mena, B., Le Moan, A., Christensen, A., Deurs, M., Mosegaard, H., Hemmer-Hansen, J., Bekkevold, D. (2020). Weak genetic structure despite strong genomic signal in lesser sandeel in the North Sea. Evolutionary Applications, 13, 376–387. doi:10.1111/eva.12875.

Jombart, T., Ahmed, I. (2011). adegenet 1.3-1: new tools for the analysis of genome-wide SNP data. Bioinformatics, 27, 3070–3071. doi:10.1093/bioinformatics/btr521

Jombart, T., Devillard, S., Balloux, F. (2010). Discriminant analysis of principal components: a new method for the analysis of genetically structured populations. BMC Genetics, 11, 94. doi:10.1186/1471-2156-11-94

Krakau, M., Jacobsen, S., Jensen, K.T., Reise, K. (2012). The cockle *Cerastoderma edule* at Northeast Atlantic shores: genetic signatures of glacial refugia. Marine Biology, 159, 221–230. doi:10.1007/s00227-011-1802-8

Lallias, D., Boudry, P., Lapegue, S., King, J.W., Beaumont, A.R. (2010). Strategies for the retention of high genetic variability in European flat oyster (*Ostrea edulis*) restoration programmes. Conservation Genetics, 11, 1899–1910. doi:10.1007/s10592-010-0081-0

Langmead, B., Trapnell, C., Pop, M., Salzberg, S.L. (2009). Ultrafast and memory-efficient alignment of short DNA sequences to the human genome. Genome Biology, 10, R5. doi.org/10.1186/gb-2009-10-3-r25

López, A., Vera, M., Planas, M., Bouza, C. (2015). Conservation genetics of threatened *Hippocampus guttulatus* in vulnerable habitats in NW Spain: temporal and spatial stability of wild populations with flexible polygamous mating system in captivity. PLoS ONE, 10, e0117538. doi.org/10.1371/journal.pone.0117538

López-Jamar, E., Cal, R.M., González, G., Hanson, R.B., Rey, J., Santiago, G., Tenore, K.R. (1992). Upwelling and outwelling effects on the benthic regime of the continental-shelf off Galicia, NW Spain. Journal of Marine Research, 50, 465–488. doi:10.1357/002224092784797584

Luttikhuizen, P.C., Campos, J., van Bleijswijk, J., Peijnenburg, K.T.C. A., van der Veer, H W. (2008). Phylogeography of the common shrimp, *Crangon crangon* (L.) across its distribution range. Molecular Phylogenetics and Evolution, 46, 1015–1030. doi:10.1016/j.ympev.2007.11.011

Maggs, C.A., Castilho, R., Foltz, D., Henzler, C., Jolly, M.T., Kelly, J., Olsen, J., Pérez, K.E., Stam, W., Väinolä, R., Viard, F., Wares, J. (2008). Evaluating signatures of glacial refugia for north atlantic benthic marine taxa. Ecology, 89, S108–S122. doi:10.1890/08-0257.1

Mahony, K.E., Lynch, S.A., Egerton, S., Cabral, S., de Montaudouin, X., Fitch A., Magalhães, L., Rocroy, M., Culloty, S.C. (2020). Mobilisation of data to stakeholder communities. Bridging the research-practice gap using a commercial shellfish species model. PLoS One, 15: e0238446. doi.org/10.1371/journal.pone.0238446

Malham, S.K., Hutchinson, T.H., Longshaw, M. (2012). A review of the biology of European cockles (Cerastoderma spp.). Journal of the Marine Biological Association of the United Kingdom, 92, 1563–1577. doi:10.1017/s0025315412000355

Maroso, F., Pérez de Gracia, C., Iglesias, D., Cao, A., Díaz, S., Villalba, A., Vera, M., Martínez, P. (2019). A useful SNP panel to distinguish two cockle species, *Cerastoderma edule* and *C. glaucum*, co-occurring in some European beds, and their putative hybrids. Genes, 10, 760. doi:10.3390/genes10100760

Marquardt, D.W. (1970). Generalized inverses, ridge regression, biased linear estimation and nonlinear estimation. Technometrics, 12, 59.

Martínez, L., Méndez, J., Insua, A., Arias-Pérez, A., Freire, R. (2013). Genetic diversity and population differentiation in the cockle *Cerastoderma edule* estimated by microsatellite markers. Helgoland Marine Research, 67, 179–189. doi:10.1007/s10152-012-0314-3

Martínez, L., Freire, R., Arias-Pérez, A., Méndez, J., Insua, A. (2015). Patterns of genetic variation across the distribution range of the cockle *Cerastoderma edule* inferred from microsatellites and mitochondrial DNA. Marine Biology, 162, 1393–1406. doi:10.1007/s00227-015-2676-y

Nielsen, E.E., Nielsen, P.H., Meldrup, D., Hansen, M.M. (2004). Genetic population structure of turbot (*Scophthalmus maximus* L.) supports the presence of multiple hybrid zones for marine fishes in the transition zone between the Baltic Sea and the North Sea. Molecular Ecology, 13, 585–595. doi.org/10.1046/j.1365-294X.2004.02097.x

Nielsen, E.E., Hemmer-Hansen, J., Larsen, P.F., Bekkevold, D. (2009). Population genomics of marine fishes: Identifying adaptive variation in space and time. Molecular Ecology, 18, 3128–3150. doi.org/10.1111/j.1365-294X.2009.04272.x

Norris, K., Bannister, R.C.A., Walker, P.W. (1998). Changes in the number of oystercatchers *Haematopus ostralegus* wintering in the Burry Inlet in relation to the biomass of cockles *Cerastoderma edule* and its commercial exploitation. Journal of Applied Ecology, 35, 75–85. doi:10.1046/j.1365-2664.1998.00279.x

Oksanen, J. (2015). Multivariate Analysis of Ecological Communities in R: vegan tutorial. Available from http://cc.oulu.fi/~jarioksa/opetus/metodi/vegantutor.pdf.

Petereit, C., Bekkevold, D., Nickel, S., Dierking, J., Hantke, H., Hahn, A., Reusch, T., Puebla, O. (2018). Population genetic structure after 125 years of stocking in sea trout (*Salmo trutta* L.). Conservation Genetics, 19, 1123–1136.

Piñeira, J., Quesada, H., Rolan-Alvarez, E., Caballero, A. (2008). Genetic discontinuity associated with an environmentally induced barrier to gene exchange in the marine snail *Littorina saxatilis*. Marine Ecology Progress Series, 357, 175–184. doi:10.3354/meps07278.

Pino-Querido, A., Álvarez-Castro, J.M., Vera, M., Pardo, B.G., Fuentes, J., Martínez, P. (2014). A molecular tool for parentage analysis in the Mediterranean mussel (*Mytilus galloprovincialis*). Aquaculture Research, 46, 1721–1735.

Raj, A., Stephens, M., Pritchard, J.K. (2014). fastSTRUCTURE: variational inference of population structure in large SNP data sets. Genetics, 197, 573–U207. doi:10.1534/genetics.114.164350

Riquet, F., Liautard-Haag, C., Woodall, L., Bouza, C., Louisy, P., Hamer, B., Otero-Ferrer, F., Aublanc, P., Béduneau, V., Briard, O., El Ayari, T., Hochscheid, S., Belkhir, K., Arnaud-Haond, S., Gagnaire, P. & Bierne, N. 2019. Parallel pattern of differentiation at a genomic island shared between clinal and mosaic hybrid zones in a complex of cryptic seahorse lineages. Evolution, 73, 817–835. doi.org/10.1111/evo.13696

Rohlf, F. (1993). NTSYS-pc. Numerical taxonomy and multivariate analysis system, Version 2.1. Setauket, New York.

Rosenberg, N.A. (2004). DISTRUCT: a program for the graphical display of population structure. Molecular Ecology Notes, 4, 137–138. doi:10.1046/j.1471-8286.2003.00566.x

Rousset, F. (1997). Genetic differentiation and estimation of gene flow from F-statistics under isolation by distance. Genetics, 145, 1219–1228.

Rousset, F. (2008). GENEPOP ’ 007: a complete re-implementation of the GENEPOP software for Windows and Linux. Molecular Ecology Resources, 8, 103–106. doi:10.1111/j.1471-8286.2007.01931.x

Rowley, A.F., Cross, M.E., Culloty, S.C., Lynch, S.A., Mackenzie, C.L., Morgan, E., O’Riordan, R.M., Robins, P.E., Smith, A.L., Thrupp, T.J., Vogan, C.L., Wootton, E.C., Malham, S.K. (2014). The potential impact of climate change on the infectious diseases of commercially important shellfish populations in the Irish Sea—a review. ICES Journal of Marine Science, 71, 741–759. doi.org/10.1093/icesjms/fst234

Scrope-Howe, S., Jones, D.A. (1986). The vertical distribution of zooplankton in the western Irish Sea. Estuarine, Coastal and Shelf Science, 22, 785–802.

Sotelo, G., Duvetorp, M., Costa, D., Panova, M., Johannesson, K., Faria, R. (2020). Phylogeographic history of flat periwinkles, *Littorina fabalis* and *L. obtusata*. BMC Evolutionary Biology, 20, 23. doi:10.1186/s12862-019-1561-6

Sotillo, M.G., Cailleau, S., Lorente, P., Levier, B., Aznar, R., Reffray, G., Amo-Baladrón, A., Chanut, J., Benkiran M., Alvarez-Fanjul, E. (2015). The MyOcean IBI Ocean Forecast and Reanalysis Systems: operational products and roadmap to the future Copernicus Service. Journal of Operational Oceanography, 8, 63–79. doi:10.1080/1755876X.2015.1014663.

Sotillo, M.G., Levier, B., Lorente, P., Guihou, K., Aznar, R. (2020). MyOcean Quality Information Document for the Atlantic - Iberian Biscay Irish-Ocean Physics Analysis and Forecasting Product (IBI_ANALYSIS_FORECAST_PHYS_005_001_b). MyOcean Technical Report (www.myocean.eu)

Sournia, A., Birrien, J.L., Camus, P., Daniel, J.Y., Jacq, E., Koutsikopoulos, C., Lecorre, P., Lecoz, J.R., Lesaos, J.P., Mariette, V., Martinjezequel, V., Moal, J., Morin, P., Poulet, S.A., Prieur, D., Samain, J.F., Slawyk, G., Videau, C. (1988). A physical, chemical and biological charaterization of the Ushant tidal front. Internationale Revue Der Gesamten Hydrobiologie, 73, 511–536.

Sromek, L., Forcioli, D., Lasota, R., Furla, P., Wolowicz, M. (2019). Next-generation phylogeography of the cockle *Cerastoderma glaucum*: Highly heterogeneous genetic differentiation in a lagoon species. Ecology and Evolution, 9, 4667–4682. doi:10.1002/ece3.5070

Teles-Machado, A., Peliz, Á., McWilliams, J.C., Couvelard, X., Ambar, I. (2016). Circulation on the Northwestern Iberian Margin: Vertical structure and seasonality of the alongshore flows. Progress in Oceanography, 140, 134–153. doi.org/10.1016/j.pocean.2015.05.021

Van Wyngaarden, M., Snelgrove, P.V.R., DiBacco, C., Hamilton, L.C., Rodriguez-Ezpeleta, N., Jeffery, N.W., Stanley, R.R.E., Bradbury, I. R. (2017). Identifying patterns of dispersal, connectivity and selection in the sea scallop, *Placopecten magellanicus*, using RADseq-derived SNPs. Evolutionary Applications, 10, 102–117. doi:10.1111/eva.12432

Vandamme, S.G., Maes, G.E., Raeymaekers, J.A.M., Cottenie, K., Imsland, A K., Hellemans, B., Lacroix, GG., Mac Aoidh, Volckaert, F A.M. (2014). Regional environmental pressure influences population differentiation in turbot (*Scophthalmus maximus*). Molecular Ecology, 23, 618–636. doi:10.1111/mec.12628

Vera, M., Carlsson, J., El Carlsson, J., Cross, T., Lynch, S., Kamermans, P., Villalba, A., Culloty, S., Martínez, P. (2016). Current genetic status, temporal stability and structure of the remnant wild European flat oyster populations: conservation and restoring implications. Marine Biology, 163, 239. doi:10.1007/s00227-016-3012-x

Vera, M., Pardo, B.G., Cao, A., Vilas, R., Fernández, C., Blanco, A., Gutiérrez, A.P., Bean, T.P., Villalba, A., Martínez, P. (2019). Signatures of selection for bonamiosis resistance in European flat oyster (*Ostrea edulis*): New genomic tools for breeding programs and management of natural resources. Evolutionary Applications, 12, 1781–1796. doi:10.1111/eva.12832

Vilas, R., Vandamme, S. G., Vera, M., Souza, C., Maes, G. E., Volckaert, F. A. M., Martínez, P. (2015). A genome scan for candidate genes involved in the adaptation of turbot (*Scophthalmus maximus*). Marine Genomics, 23, 77–86. doi.org/10.1016/j.margen.2015.04.011

Villalba, A., Iglesias, D., Ramilo, A., Darriba, S., Parada, J.M., No, E., Abollo, E., Molares, J., Carballal, M.J. (2014). Cockle *Cerastoderma edule* fishery collapse in the Ría de Arousa (Galicia, NW Spain) associated with the protistan parasite *Marteilia cochillia*. Diseases of Aquatic Organisms, 109, 55–80. doi:10.3354/dao02723.

Weir, B.S., Cockerham, C.C. (1984). Estimating f-statistics for the analysis of population-structure. Evolution, 38, 1358–1370.

Xie, C., Mao, X., Huang, J., Ding, Y., Wu, J., Dong, S., Kong, L., Gao, G., Li, C. Y., Wei, L. (2011). KOBAS 2.0: a web server for annotation and identification of enriched pathways and diseases. Nucleic Acids Research, 39(Web Server issue), W316–W322. doi:10.1093/nar/gkr483.

Xuereb, A., Kimber, C. M., Curtis, J.M.R., Bernatchez, L., Fortin, M. J. (2018). Putatively adaptive genetic variation in the giant California sea cucumber (*Parastichopus californicus*) as revealed by environmental association analysis of restriction-site associated DNA sequencing data. Molecular Ecology, 27, 5035–5048. doi:10.1111/mec.14942.

Zhao, Y., Peng, W., Guo, H., Chen, B., Zhou, Z., Xu, J., Zhang, D., Xu, P. (2018). Population Genomics reveals genetic divergence and adaptive differentiation of Chinese sea bass (*Lateolabrax maculatus*). Marine Biotechnoly, 20, 45–59. doi:10.1007/s10126-017-9786-0.

