## Supplementary Figures for "Biotic and abiotic factors shaping the genome of cockle (*Cerastoderma edule*) in the Northeast Atlantic: a baseline for sustainable management of its wild resources"

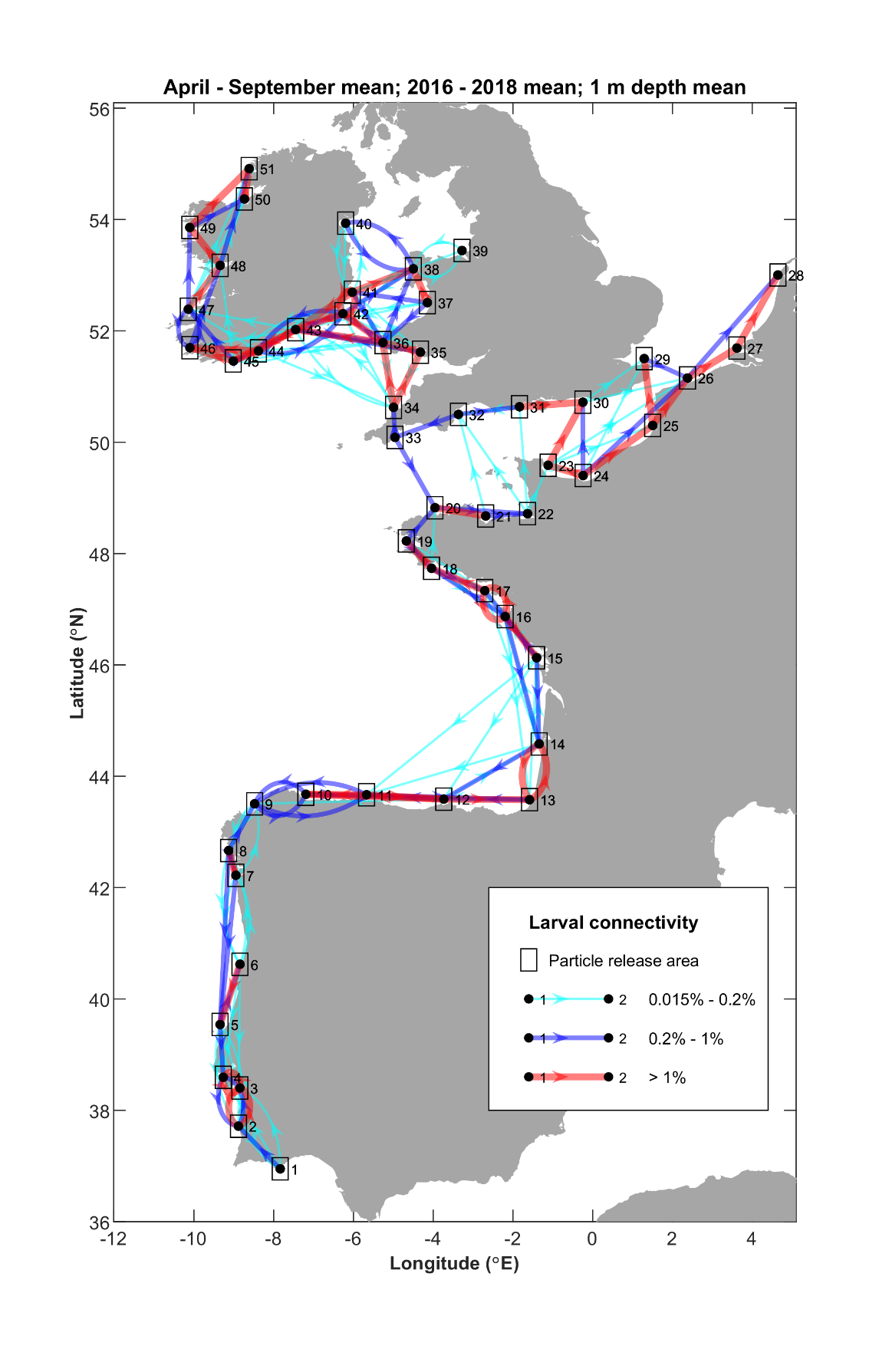

**Supplementary Figure 1**: Mean larval connectivity pathways for April to September from 2016 – 2018 releases 1 m depth. The direction of the arrows indicates the direction of larval transport and the colour and thickness of the connection displays the strength of the connection.

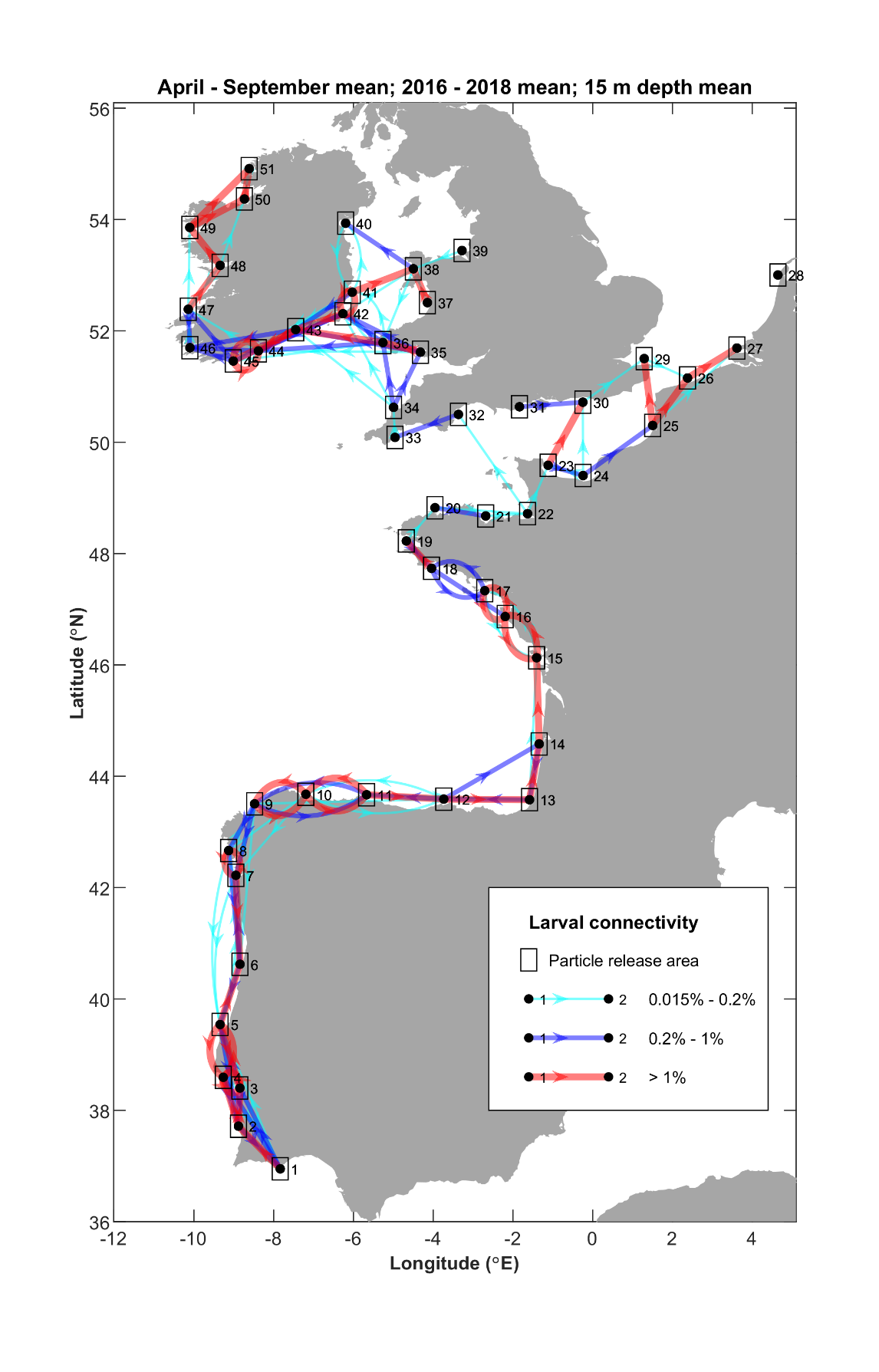

**Supplementary Figure 2**: Mean larval connectivity pathways for April to September from 2016 – 2018 releases 15 m depth. The direction of the arrows indicates the direction of larval transport and the colour and thickness of the connection displays the strength of the connection.

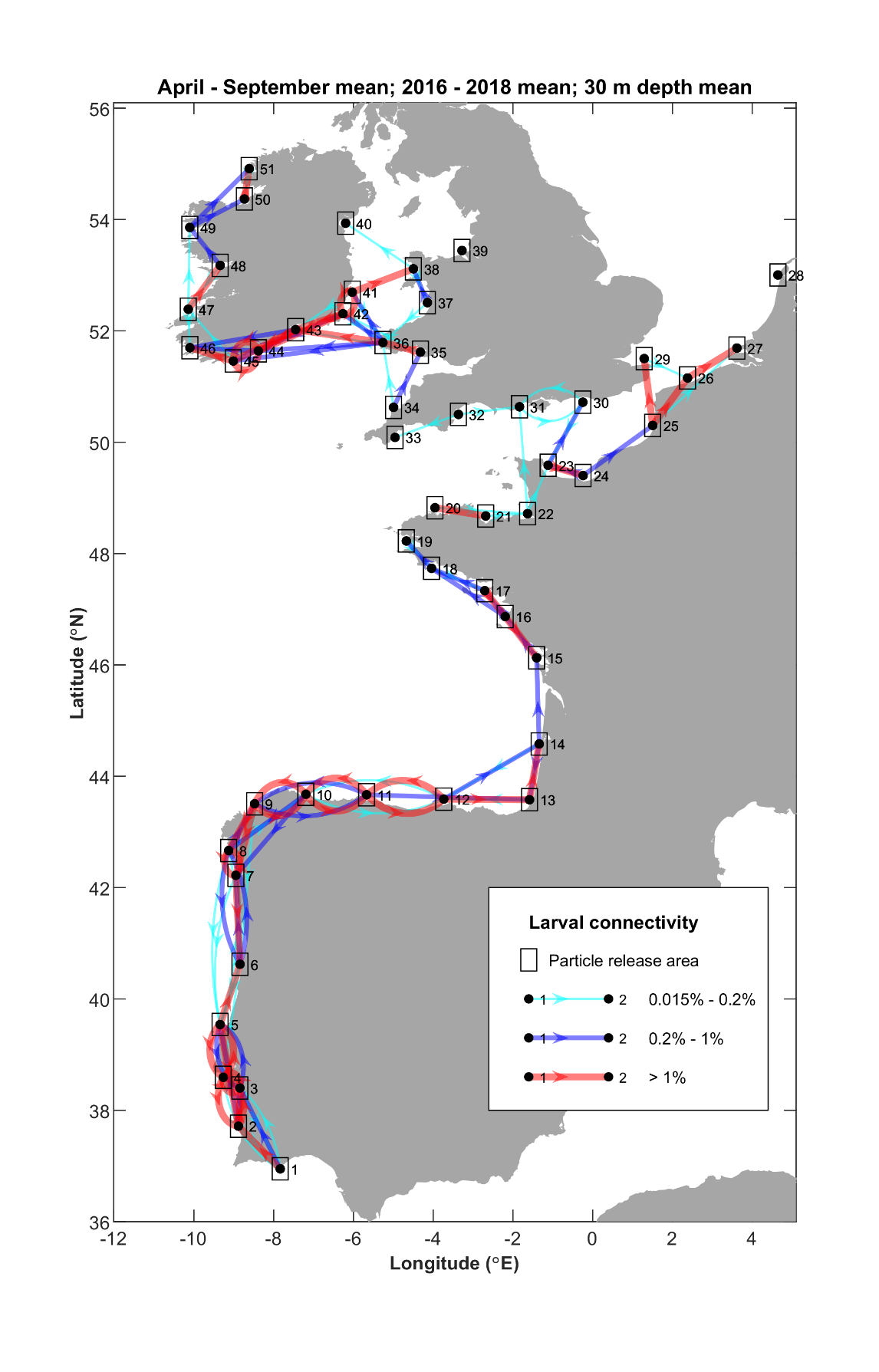

**Supplementary Figure 3**: Mean larval connectivity pathways for April to September from 2016 – 2018 releases 30 m depth. The direction of the arrows indicates the direction of larval transport and the colour and thickness of the connection displays the strength of the connection.

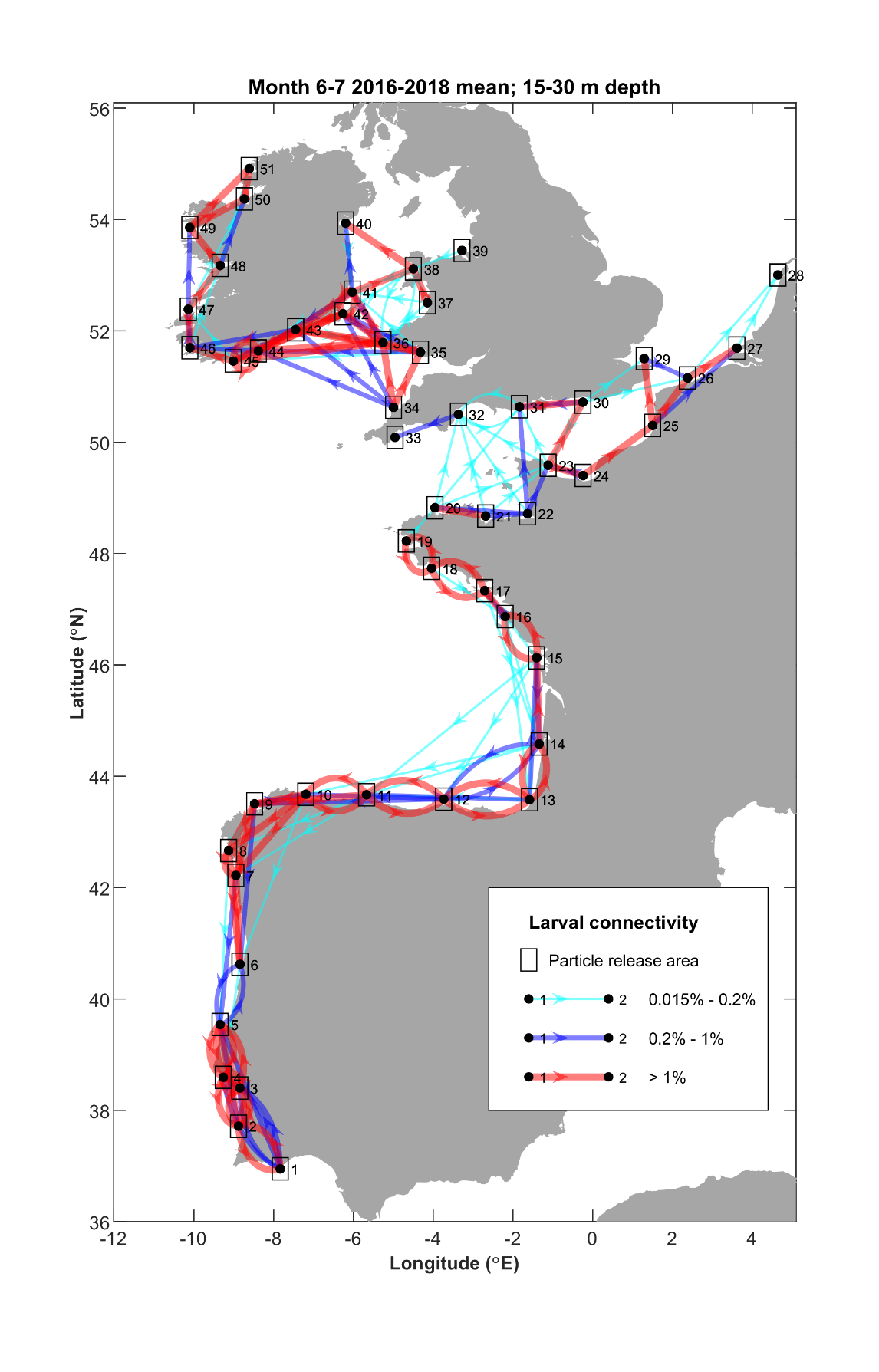

**Supplementary Figure 4:** Mean larval connectivity pathways for June to July from 2016 – 2018 releases for 15 and 30 m depth. The direction of the arrows indicates the direction of larval transport and the colour and thickness of the connection displays the strength of the connection.

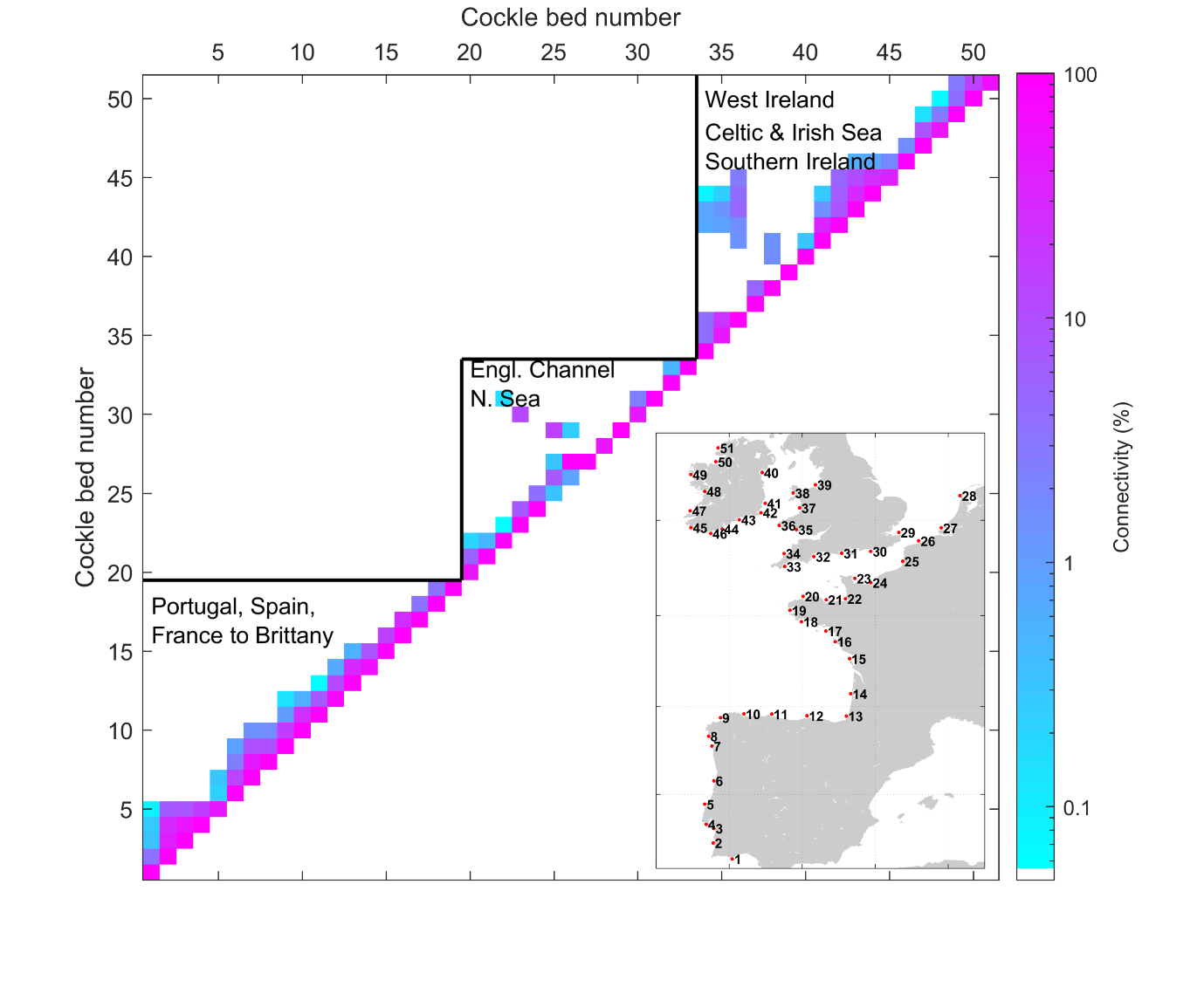

**Supplementary Figure 5:** Mean larval connectivity matrix for June to July from 2016 – 2018 for 15 and 30 m depth releases. The strength of the connectivity between two sites is shaded and the location of each site can be seen in the map in the right-hand bottom corner. Distinct regions are indicated by the black lines.
